# A functional investigation of antibody Fc-FcRn variant binding guided by *in silico* free energy perturbation methods

**DOI:** 10.64898/2026.04.28.721095

**Authors:** Jared M. Sampson, Alina P. Sergeeva, Tianyang Gao, Young Do Kwon, Eswar Reddem, Fabiana A. Bahna, Seetha M. Mannepalli, Baoshan Zhang, Peter D. Kwong, Lawrence Shapiro, Barry Honig, Richard A. Friesner

**Affiliations:** Department of Biochemistry and Molecular Biophysics, Columbia University, New York, NY, USA, 10027; Department of Chemistry, Columbia University, New York, NY, USA, 10027; Life Sciences Software, Schrödinger, Inc., New York, NY, USA, 10036; Department of Systems Biology, Columbia University, New York, NY, USA, 10032; Vaccine Research Center, National Institute of Allergy and Infectious Diseases, National Institutes of Health, Bethesda, MD, USA, 20892; Zuckerman Mind Brain Behavior Institute, Columbia University, New York, NY, USA, 10027; Aaron Diamond AIDS Research Center, Columbia University, New York, NY, USA, 10032; Department of Medicine, Columbia University, New York, NY, USA, 10032

**Author notes:** Corresponding author: Email addresses (Barry Honig), (Richard A. Friesner).

**Keywords:** antibody-receptor interactions, binding affinity prediction, free-energy methods

## Abstract

Accurate calculation of energy changes upon mutation is a key requirement for the effective use of computational methods in protein design. In this study, we applied free energy perturbation (FEP) calculations to predict the effects of mutations on the binding free energy between the immunoglobulin subtype G (IgG) antibody fragment-crystallizable (Fc) region and the neonatal Fc receptor (FcRn), an interaction that is primarily responsible for antibody half-life. We assembled an extensive experimental dataset of Fc-FcRn binding affinities for wild-type (*wt*) and mutant complexes, including values from literature and from newly measured results. Starting from a crystal structure of the M252Y/S254T/T256E (“YTE”) Fc variant bound to FcRn, we prepared all-atom models of human IgG1-subtype *wt* and YTE variant Fc-FcRn complexes, adding explicit hydrogens and assigning protonation states for key ionizable residues. Initial results using standard FEP protocols to compute relative binding free energies were promising but exhibited multiple outliers. By accounting for coupling effects for FEP mutations near key histidine residues, we improved the results for several outliers, suggesting such coupling as an important approach for pH-sensitive systems. Further, upon determining new crystal structures of four Fc variants at multiple pH values, we observed subtle conformational changes in unbound Fc; by accounting for these conformational changes in FEP calculations, we additionally improved agreement with experiment. The detailed structural and energetic analyses of the Fc-FcRn system we present here thus provide an accurate energy-calculation framework to enable rational *in silico* design of novel Fc variants.

**Significance:** The ability to determine changes in binding affinity upon mutation is critical to structure-based protein design. In this study, we demonstrate a successful computational approach using free energy perturbation (FEP) calculations on the antibody Fc-FcRn complex, a medically relevant system with implications for both therapeutic and prophylactic antibody use. Our successful calculation of accurate binding energies across a wide range of cases speaks to the power of the FEP methodology in navigating the free energy landscapes of dynamic molecular complexes. Furthermore, we show that accurate Fc-FcRn affinity calculations required careful consideration of conformational flexibility between bound and unbound states, contributing to our functional understanding of a system that will be important for future rational antibody-design efforts.

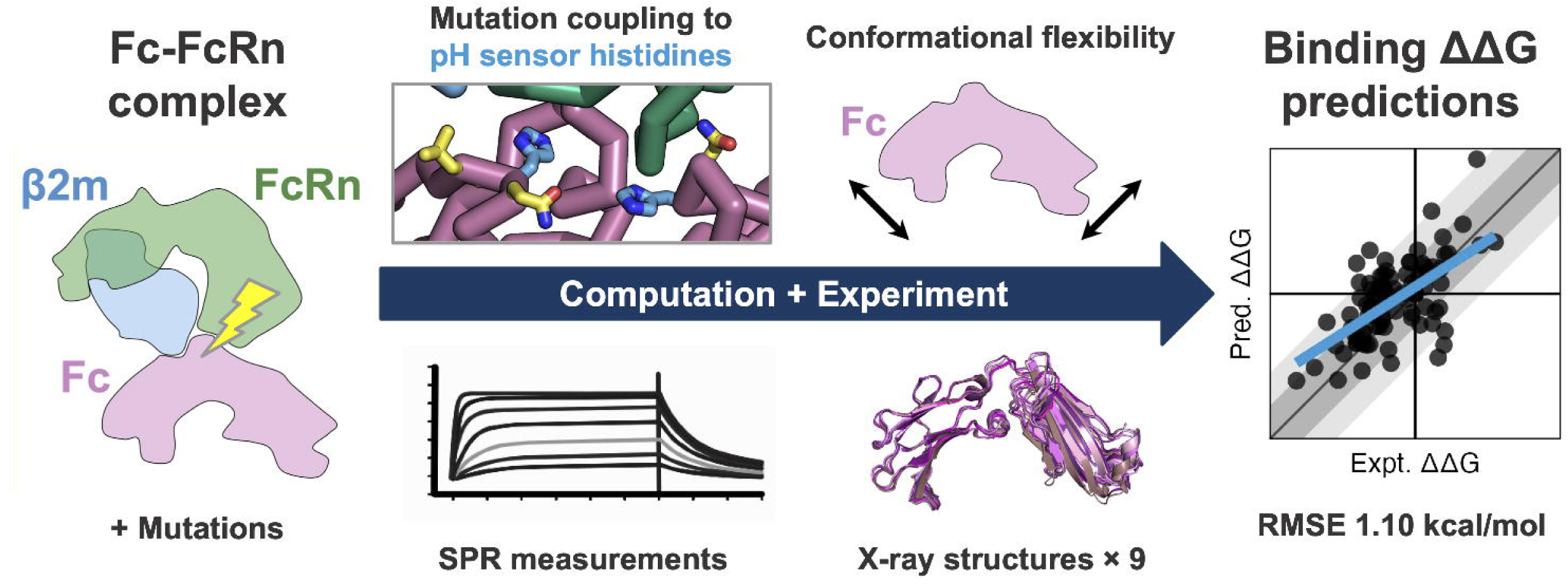

## Introduction

The neonatal Fc receptor (FcRn) plays a critical role in maintaining the circulating immunoglobulin subtype G antibody (IgG Ab) concentration in humans and other mammals. This homeostatic regulation is accomplished via a recycling mechanism in which, after nonspecific fluid-phase uptake of the antibody by pinocytosis, FcRn binds to the Ab fragment crystallizable (Fc) region in the acidified early endosome [1]. The IgG-FcRn complex is transported back to the cell surface, where, due to reduced binding affinity at serum pH, the complex dissociates, returning the antibody to circulation and freeing the receptor for reuse [1]. This pH-dependent binding to FcRn increases the serum half-life of IgG antibodies to approximately three weeks, from 5-7 days for other antibody subtypes [2]. In addition to controlling IgG antibody concentration, FcRn has a number of other functions, including transport of IgG from maternal to fetal circulation in the placenta, and facilitating IgG absorption from mother’s milk by the infant gastrointestinal epithelium, the function from which its name is derived [1, 3, 4, 5].

Over the past few decades, numerous attempts have sought to engineer the Fc region to improve IgG half-life by increasing binding affinity to FcRn in the endosome [6, 7, 8, 9, 10, 11, 12, 13, 14, 15]. Currently available modified Fc regions have been shown to increase IgG half-life by approximately 2- to 6-fold, but further optimization leading to larger increases in half-life could potentially enhance the utility of antibodies in prophylactic and other long-term applications. While substantial progress has been made in producing Fc variants with altered binding affinities, the current best-performing Fcs have all been identified using experimental display and sorting techniques and may be suboptimal with regard to the binding pH-dependence that would maximize antibody half-life. Ideally, this would involve very high affinity binding to FcRn at acidic pH (∼6.0) in the early endosome and complete release back into serum at pH 7.4. The challenge, then, is to design antibodies whose FcRn-binding properties vary significantly over this pH range. However, accounting for protonation state and pH sensitivity presents a substantial obstacle to structure-based rational design efforts, where current computational screening methods are generally empirically fit or trained without consideration of these features.

We have investigated the utility of computational methods, specifically free energy perturbation (FEP) calculations, to design an Fc region that can substantially extend antibody half-life beyond what is obtainable with currently available variants. FEP calculations consist of molecular dynamics (MD) simulations with all-atom explicit solvent and simulate an alchemical transition between two closely related molecular species—in the present case, two amino acid sidechains. We use an implementation of the FEP methodology called FEP+ (Schrödinger, Inc.), which has demonstrated success in the prediction of free energy changes for a range of systems and perturbation types, including residue mutation in protein-protein binding, as well as other applications such as small molecule ligand-receptor binding, protein stability, and protein residue pKas [16, 17, 18, 19, 20, 21, 22].

In the study reported here, we carried out an extensive FEP-based investigation of the effect of protein residue mutation on Fc-FcRn binding. We utilized the high-resolution (2.4 Å) structure of the human Fc-FcRn complex (PDB 6WNA) as the starting point for FEP calculations [23], and supplemented the available structural information from the literature with new crystal structures of unbound apo Fc variants, crystallized over a range of pH values. We employed validated simulation and analysis methods for computing pKa values of key ionizable residues from both the antibody and the receptor, as well as the effects of those pKas on binding affinities [19, 24, 22], utilizing as the energy model the most advanced version of the protein molecular mechanics force field available for FEP+ simulations, OPLS5, which incorporates explicit polarization for a key subset of residues [25].

We evaluated the agreement between theory and experiment by comparing FEP-calculated and experimental relative binding free energies (ΔΔ*G*s) for a large dataset of Fc and FcRn variants, augmenting a curated set of Fc variant cases from literature with a set of newly measured binding affinities, including data for both Fc and FcRn variants. We improved an FEP protocol for calculating the pH-specific effect of mutations on protein-protein binding, as previously proposed [22], via an enhanced treatment of a small subset of mutations which, by virtue of their proximity, are strongly coupled to the protonation states of two key histidine residues that are the putative pH sensors in the system. Simple empirical criteria were used to identify a minimal set of such residues, thus ensuring limited impact on overall computational cost.

When this updated methodology was applied to the entire dataset described above, only a small number of significant outliers (cases where predicted binding ΔΔ*G* differs by more than 2 kcal/mol from experiment) remained. The most likely explanation for these outliers involved mutation-induced conformational changes of either the Fc-FcRn complex, or the isolated Fc. We explored these possibilities by updating our FEP calculations to take the new apo Fc structural information into account. The results obtained here suggest that a systematic effort integrating FEP calculations with structural biology can be used to achieve a robust understanding of the free energy landscape.

## Results

### Preparation of pH-specific all-atom structural models

Using all-atom complex models of wild-type (*wt*) and YTE Fc bound to FcRn prepared as described in Materials and Methods, we calculated pKa values for key ionizable amino acid sidechains at the binding interface using the FEP-based method described by Coskun and coworkers [19]. These calculations were run using either OPLS4 or OPLS5 force fields, with the latter modeling explicit polarization of key groups, including aromatic and carboxylate amino acid sidechains [25]. Table 1 presents computed pKa values for both the *wt* and YTE Fc-FcRn complex models. Based on these pKa values and the experimental pH of 6.0, structural models were updated to include the protonation states of the interfacial ionizable residues indicated in the last column of the table. The final, pH-specific all-atom structural model for the *wt* Fc-FcRn complex is presented in Figure 1.

**Figure 1.**
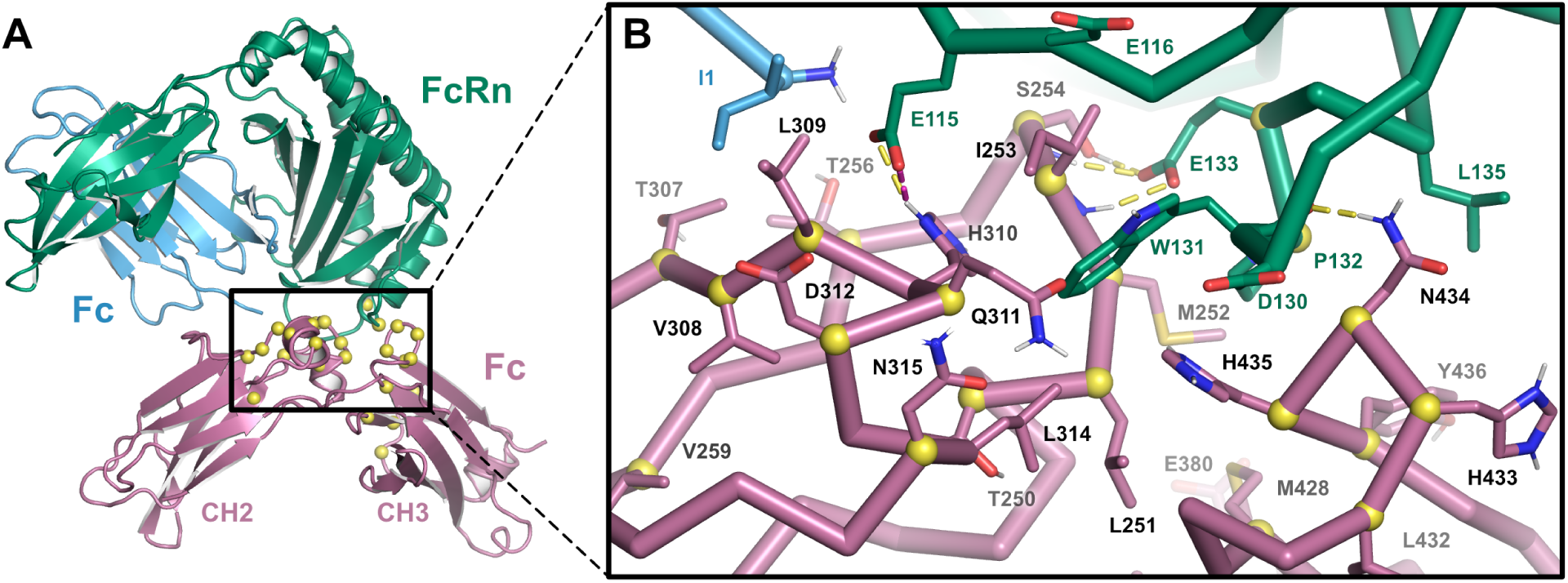
All-atom structural model of the Fc-FcRn complex used for FEP calculations. (A) Ribbon representation of the structural model of the *wt* Fc-FcRn complex prepared from the 6WNA crystal structure, with the FcRn α chain shown in green, β2 microglobulin in light blue, and Fc in violet. Sites investigated using FEP are indicated by yellow spheres. (B) Close up view of the binding interface in the same orientation as indicated by the box in (A). Protein backbone is shown as cylindrical ribbons, key Fc and FcRn/β2M residues are labeled, and *trans* hydrogen bonds and salt bridge interactions between the antibody and receptor are shown as dashed lines.

**Table 1.** FEP-calculated bound-state pKa values for interface residues in the 6WNA-derived models using OPLS4 or OPLS5.

| Chain | Residue | OPLS4 |  | OPLS5 |  | Modeled<sup>a</sup> |
| --- | --- | --- | --- | --- | --- | --- |
|  |  | WT | YTE | WT | YTE |  |
| Fc | T256 / E256 | – | 4.30 | – | 4.42 | GLU |
| Fc | H285 | 6.58 | 6.55 | 6.68 | 6.63 | HIP |
| Fc | H310 | 11.33 | 11.67 | 10.22 | 10.35 | HIP |
| Fc | D312 | 2.88 | 2.84 | 2.92 | 3.11 | ASP |
| Fc | E380 | 4.40 | 4.05 | 4.78 | 4.37 | GLU |
| Fc | E430 | 5.56 | 5.86 | 6.13 | 6.47 | GLU |
| Fc | H433 | 7.54 | 7.45 | 7.49 | 7.42 | HIP |
| Fc | H435 | 6.17 | 5.70 | 6.96 | 6.42 | HIP |
| FcRn | E115 | -0.07 | 0.91 | -0.22 | 1.79 | GLU |
| FcRn | E116 | 2.45 | 2.54 | 3.19 | 3.28 | GLU |
| FcRn | D130 | 4.05 | 4.00 | 4.12 | 3.93 | ASP |
| FcRn | E133 | 2.04 | 3.30 | 2.63 | 2.82 | GLU |

<sup>a</sup> Protonation states selected for all-atom models used in retrospective FEP calculations.

### Mutational binding affinity dataset curation

We assembled an extensive set of published binding affinities (*K*_D_) for *wt* and variant Fc-FcRn binding interactions at endosomal pH. The variant Fcs included single, double, and triple-mutants relative to either *wt* or M252Y/S254T/T256E (YTE) triple-mutant backgrounds [6, 7, 8, 9, 10, 11, 12, 13, 14, 15]. This dataset included 89 *K*_D_ data points comprising 52 unique variants, and spanned a binding ΔΔ*G* range from -3.59 to +0.77 kcal/mol relative to *wt* Fc-FcRn after converting *K*_D_ values to ΔΔ*G* and averaging across sources. Due to emphasis in the literature on improving binding between Fc and FcRn, most (93%) of these published variant measurements were favorable (with negative ΔΔ*G*, and therefore higher binding affinity) compared to *wt*, with only 5 unfavorable mutation cases.

In an effort to provide additional validation data and broaden our understanding of specific structural regions of the Fc-FcRn binding interface, we performed additional SPR binding affinity measurements for 45 Fc and FcRn variants using anti-HIV-1 monoclonal Ab VRC01 as the *wt* antibody [26]. SPR traces and analyses are included in Supplemental Figure 1, and the new binding data are summarized in Supplemental Table 1. These new measurements included 10 single amino acid substitutions in the YTE Fc background, and 5 FcRn variants binding to *wt* Fc. Altogether the combined dataset contains 134 measurements of 85 unique variants, with ΔΔ*G* values ranging from -3.59 to +2.41 kcal/mol, and are summarized in Table 2. Further analysis of the experimental dataset is shown in Supplemental Figure 2, and the entire combined dataset is included with this manuscript as Supplementary Dataset 1.

**Table 2.** Summary of the experimental datasets used for FEP calculations.

| Subset | Number of Variants<sup>a</sup> | | | | Experimental $\Delta\Delta G$<sup>b</sup> | |
| --- | --- | --- | --- | --- | --- | --- |
|  | Single | Double | Triple | Total | Min | Max |
| WT Fc | 34 | 18 | 11 | 63 | -3.59 | 2.41 |
| YTE Fc | 11 | 3 | 2 | 16 | -2.40 | 0.53 |
| WT FcRn | 5 | 0 | 0 | 5 | 0.15 | 1.86 |
| Total | 50 | 21 | 13 | 84 | -3.59 | 2.41 |

<sup>a</sup> The number of single, double, and triple mutation variants, and total number of variants in each subset are shown.
<sup>b</sup> Experimental binding $\Delta\Delta G$ range relative to the *wt* complex in kcal/mol.

### Retrospective FEP results

We performed 10 ns Protein FEP binding calculations for each Fc or FcRn variant from the experimental dataset with three or fewer mutations relative to either the *wt* or YTE starting models. The variants were divided into three data subsets based on which starting model was used (*wt* or YTE) and which protein (Fc or FcRn) was mutated. The simulations for each variant used the starting model requiring the fewest mutations. For example, the Fc variant M252Y/S254T was accessible as a double mutant from *wt* Fc, and as a single E256T reversion mutant from YTE, so this data point was included in the YTE Fc subset. Final retrospective FEP results for the three data subsets are shown in Figure 2, reflecting binding ΔΔ*G* predictions after the iterative process described in the remainder of the text. In the first stage of this process, ΔΔ*G* results obtained after applying the Protein FEP Groups treatment at pH 6.0 were reported as binding ΔΔ*G* values relative to the *wt* Fc-FcRn starting model, which was also included as a triple reversion mutation variant in the YTE Fc subset. The Protein FEP Groups treatment, originally presented in [22] and described in detail in Supplementary Methods, accounts for the effects of pH in a robust way, effectively weighting the contributions of alternate protonation microstates in the unbound and bound forms. Retrospective FEP results for the full dataset at various stages of our analysis are presented in Supplemental Figure 3, with the above-described preliminary results shown in Supplemental Figure 3A.

**Figure 2.**
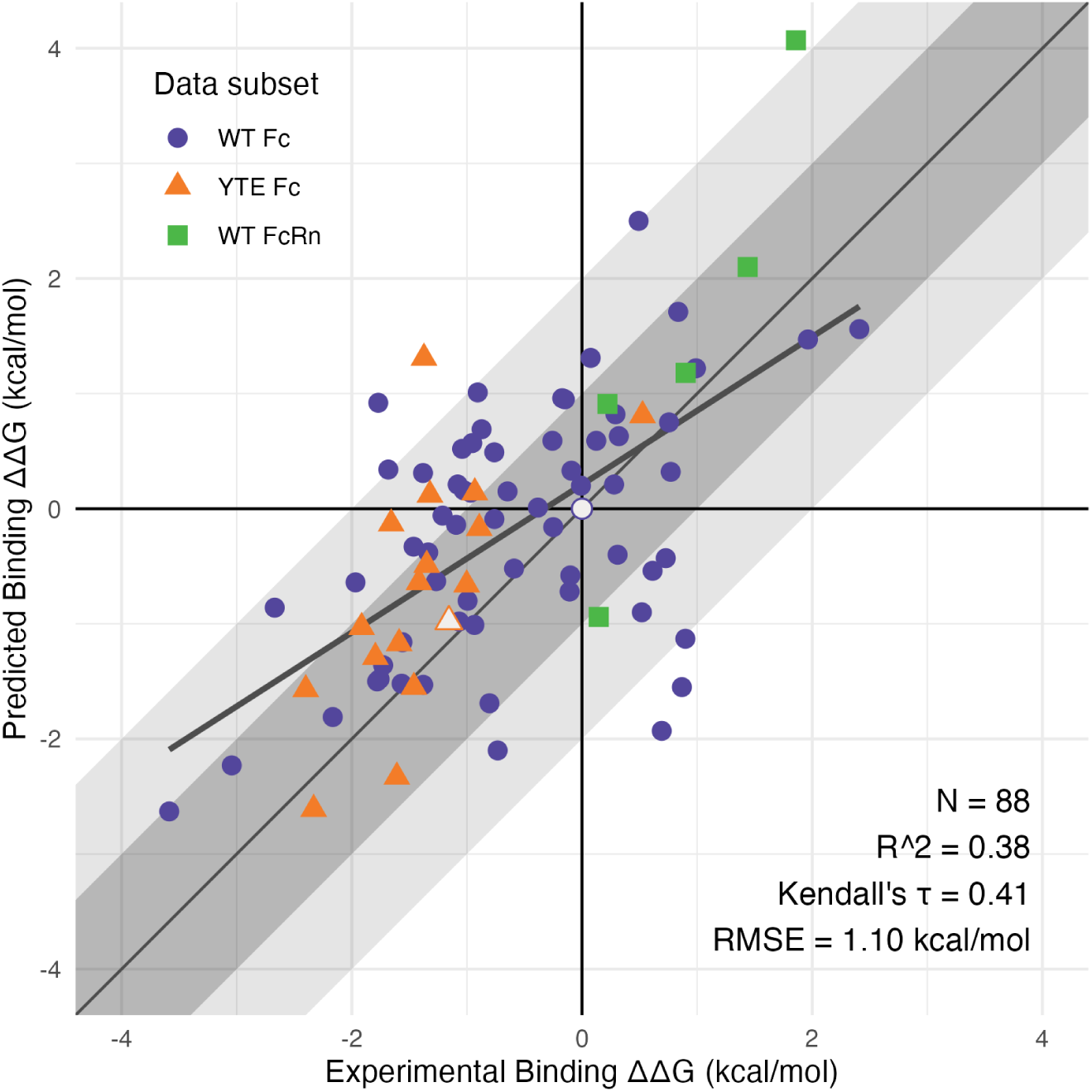
Final retrospective FEP results, showing correlation of FEP-predicted vs. experimental binding ΔΔ*G* values relative to *wt* Fc-FcRn. Point shape and color indicate which model (*wt* or YTE) was used, and which protein (Fc or FcRn) was mutated. White points indicate the results for *wt* and YTE starting models. Overall statistics are shown for the full merged dataset.

### Accounting for the effects of mutations on the ionization states of key histidine residues

With proper weighting of the contributions of protonation microstates for mutated residues accomplished via Protein FEP Groups treatment, we had addressed the most obvious requirements in the computation of the effects of mutation on binding pH dependence. However, in cases where the mutated residue was in close proximity to ionizable residues whose pKa values, and therefore protonation states, changed substantially as a result of the binding process, a more complex treatment was needed.

For variants with mutations near key Fc ionizable residues H310 and H435, we hypothesized that interactions with the mutated sites could alter the His residue pKas, and therefore affect the overall binding affinity of the mutant complex as a function of pH. From a statistical mechanical perspective, treating the sites as coupled required the calculation of binding Δ*G* values for all possible combinations of microstates, as described in the Supplementary Methods. Practically, this corresponded to sequential multiple mutation calculations such that the FEP perturbation map—a network graph in which unique protein states are represented as nodes and FEP perturbations between them as edges—included all possible microstates of the mutated residue as well as the two His pH sensors, where the total number of microstates for a given variant was the product of the individual numbers of microstates possible for each coupled titratable site.

We investigated seven cases this way, namely Fc mutations L251N, M252R, M252H, I253M, H435K, H435Y, and L309D/Q311H/N434S (“DHS”). These cases were selected as described in Materials and Methods based on empirical criteria for solvent accessibility and distance from the ionizable nitrogen atoms in the H310/H435 sidechain imidazole groups. The different identities of the mutated residues at these coupled sites led to varying levels of complexity in the setup of the FEP calculations. For example, in the context of L251N, I253M, or M252R, 9 microstates were required, enumerating all combinations of H310 and H435 protonation states (HID/HIE/HIP), which were included in both the *wt* and mutant contexts. Each distinct combined microstate was represented as a node in the FEP perturbation map. The M252H mutation introduced a third His residue, therefore requiring 27 nodes in the fully coupled FEP map corresponding to that variant. The H435K and H435Y mutations each eliminated one of the coupled His sites in question (H435), thus reducing the number of microstates in the mutant context, though in the case of H435K this was replaced with a titratable lysine sidechain. The most complex and computationally expensive variant treated in this manner was DHS, with 54 possible microstates for the L309D/Q311H/N434S triple mutant alone, with additional nodes representing microstates for the intermediate single and double mutants. Accordingly, to reduce computational costs, an intermediate L309D/Q311H starting model was used for the DHS simulations. This reduced the number of perturbations required to sample all the microstates, but relied on the assumption that the neutral N434S mutation, with a Cβ-Cβ distance of about 6 Å to the closest of the coupled titratable sites (H435), did not substantially affect the protonation states of any of these titratable residues. The FEP perturbation map prepared for the DHS variant alone initially contained over 350 perturbation edges. However, the edges in the automatically generated map were highly redundant, allowing the systematic removal of approximately half of the edges before submitting the calculation, leaving 109 nodes and 171 edges, which was sufficient to fully connect the graph, with each node included in a closed cycle containing 4 or fewer nodes. An illustrative example of these FEP graphs is presented in Supplemental Figure 4.

After performing the additional calculations to account for the coupling of these seven variants to H310 and H435, we observed a range of effects. While the effect for the L251N was negligible and the prediction for M252H resulted in a higher absolute error by 0.5 kcal/mol, M252R and I253M showed modest improvements relative to uncoupled calculations, with absolute errors decreasing by about 0.2 and 0.6 kcal/mol, respectively. The H435K and H435Y variants also demonstrated some improvement when the effects of coupling to H310 were included, with absolute errors reduced by 0.7 and 0.1 kcal/mol, respectively. The DHS variant exhibited the largest coupling effect, with a reduction in absolute error of about 1.8 kcal/mol. A summary of the changes in predicted binding ΔΔ*G* for these variants can be found in Supplemental Table 2, and the direction and magnitude of the ΔΔ*G* shifts are shown as vertical tails in the right panel of Supplemental Figure 3B.

When binding ΔΔ*G* values from these His-coupled cases were merged into the full dataset and compared to experiment, the resulting *R*^2^ of 0.34 and RMSE of 1.15 kcal/mol were comparable to previously reported protein binding FEP results [17, 18, 22]. However, there remained a small number of significant outliers, including a cluster of cases with mutations on the Fc 252-256 loop exhibiting similar “overbound” behavior—with predicted ΔΔ*G* more favorable than the experimental value, as seen at the lower right of Supplemental Figure 3B—which we decided to investigate via crystallography and further FEP calculations.

### Conformational heterogeneity

For many residues in an FEP simulation, the correct, lowest-energy region of phase space is likely to be already present in, or readily accessible from, the starting conformations of the *wt* and mutant proteins. However, for a small (but potentially significant) subset of residues, the lowest free energy structure in the context of either the complex, or the isolated protein in solution, may not be properly sampled in the course of a typical-duration FEP simulation. Ideally, in the absence of variant-specific experimental structures, one would use protein structure prediction and refinement methodology to identify the most likely starting conformation for every residue. In practice, an approach of this type is not yet feasibly within the capabilities of current refinement algorithms. Consequently, we adopted an iterative methodology in which we initially relied on the FEP simulations to sample the mutant conformational phase space, identified outliers in binding affinity prediction compared to experiment, and investigated the outliers via both computational and experimental means.

### Conformational flexibility in Fc monomer structures

We hypothesized that the observed errors for the overbound subset of outlier cases arose from inadequate representation of the unbound Fc protein due to the existence of a lower-energy conformation that was not sampled in FEP simulations. To test this hypothesis, we determined crystal structures of *wt* Fc and three Fc variants associated with outlier cases (M252H, M252R, I253M) in their unbound apo forms, each at multiple pH values (see Materials and Methods for details on crystallization conditions). Crystallographic data collection and refinement statistics for the 9 structures are presented in Supplemental Table 3.

All apo Fc crystal structures were solved as Fc homodimers; however, monomer representations are shown in Figure 3A to illustrate the overall conformational changes affecting CH2-CH3 interdomain orientation. With CH2 domains aligned, the CH3 domains exhibited translational displacements varying from 0.1 to 6.0 Å, and angular shifts ranging from 0.5° to 20.7° relative to three different reference structures (Supplemental Table 4).

**Figure 3.**
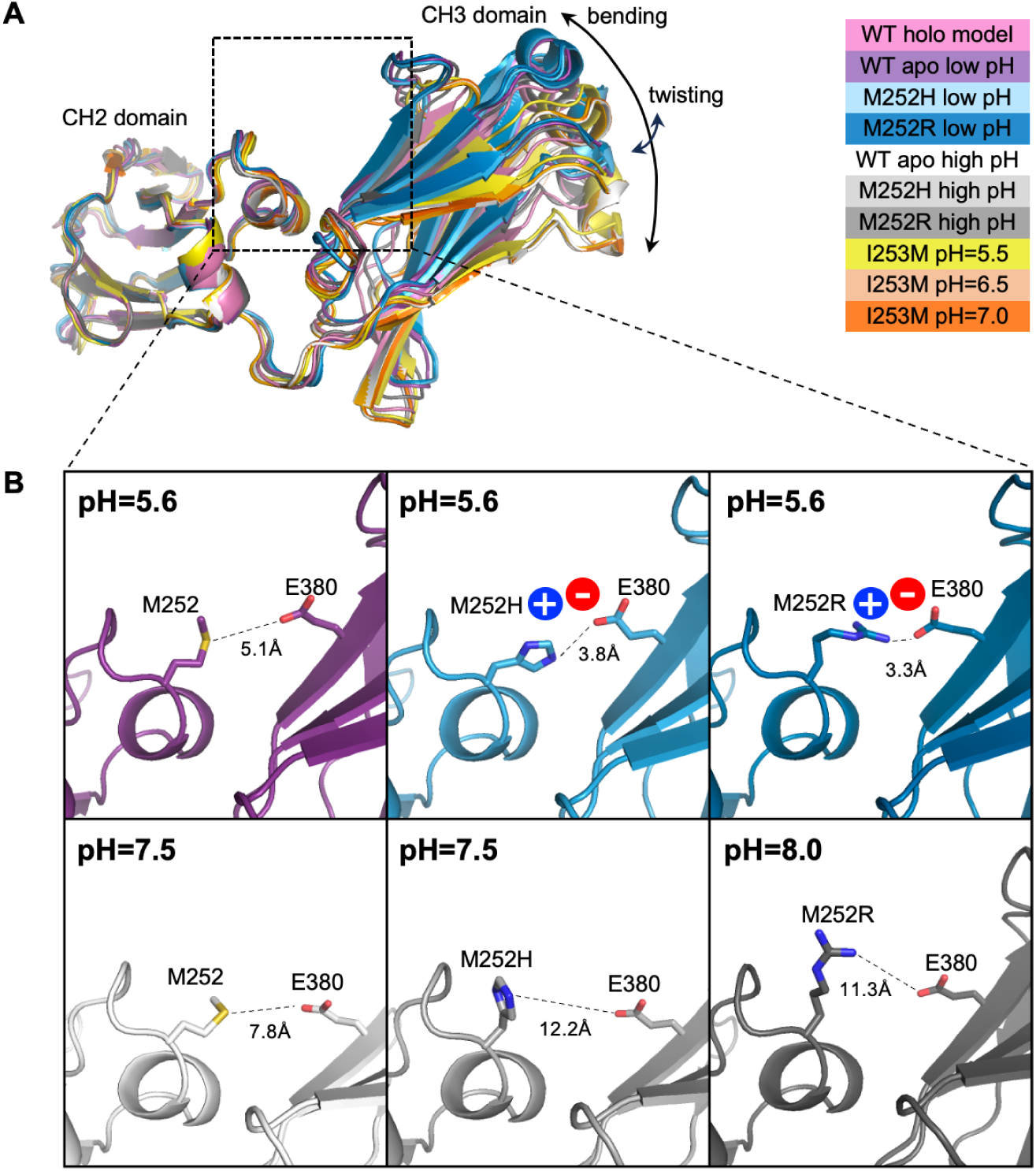
Conformational changes in the Fc protein induced by mutation or pH variation. (A) Ribbon representation of Fc monomer structures crystallized in this study, superimposed on the CH2 domain of the *wt* Fc in the bound state (holo, pink). A detailed structural comparison, including the quantified angle and displacement of the CH3 domain, is provided in Supplemental Table 4. (B) Close-up views of the Fc region highlighted by the dashed box in (A). Residues contributing to enhanced electrostatic interactions in the mutated environment are shown as sticks. Distances between the closest heavy atoms of Fc residues 252 and 380 are shown as dashed lines, with salt bridge formation indicated by circles labeled with residue charges. Structures are color-coded according to the legend in (A). At high pH (bottom row), the distance between residues 252 and 380 is greater compared to low pH (top row).

Normal mode analysis was performed on Fc chains from the original holo model, each new unbound Fc structure, and a dimer Fc model prepared from a lower resolution YTE Fc-FcRn structure (PDB 4N0U). In all structures (both monomers and the dimer), the lowest-frequency mode corresponded to a hinge-like “bending” motion, analogous to the symmetric in-plane bending vibrational mode of a water molecule. The second-lowest-frequency mode represented an out-of-plane “twisting” motion. Linear combinations of these two lowest-frequency modes could describe transitions between all observed structures. The bending mode predominantly contributed to the transition between holo and apo *wt* structures, while the twisting mode appeared to play a role in the transition between low and high pH apo *wt* structures. Regardless of the oligomeric state (monomeric or dimeric Fc), the two lowest-frequency modes remained consistent across both *wt* and mutant proteins and accounted for the structural variability observed in crystallographic data. The observed conformations in the new apo Fc structures corresponding to these domain-level “bending” and “twisting” motions of the Fc molecules are depicted in Figure 4.

**Figure 4.**
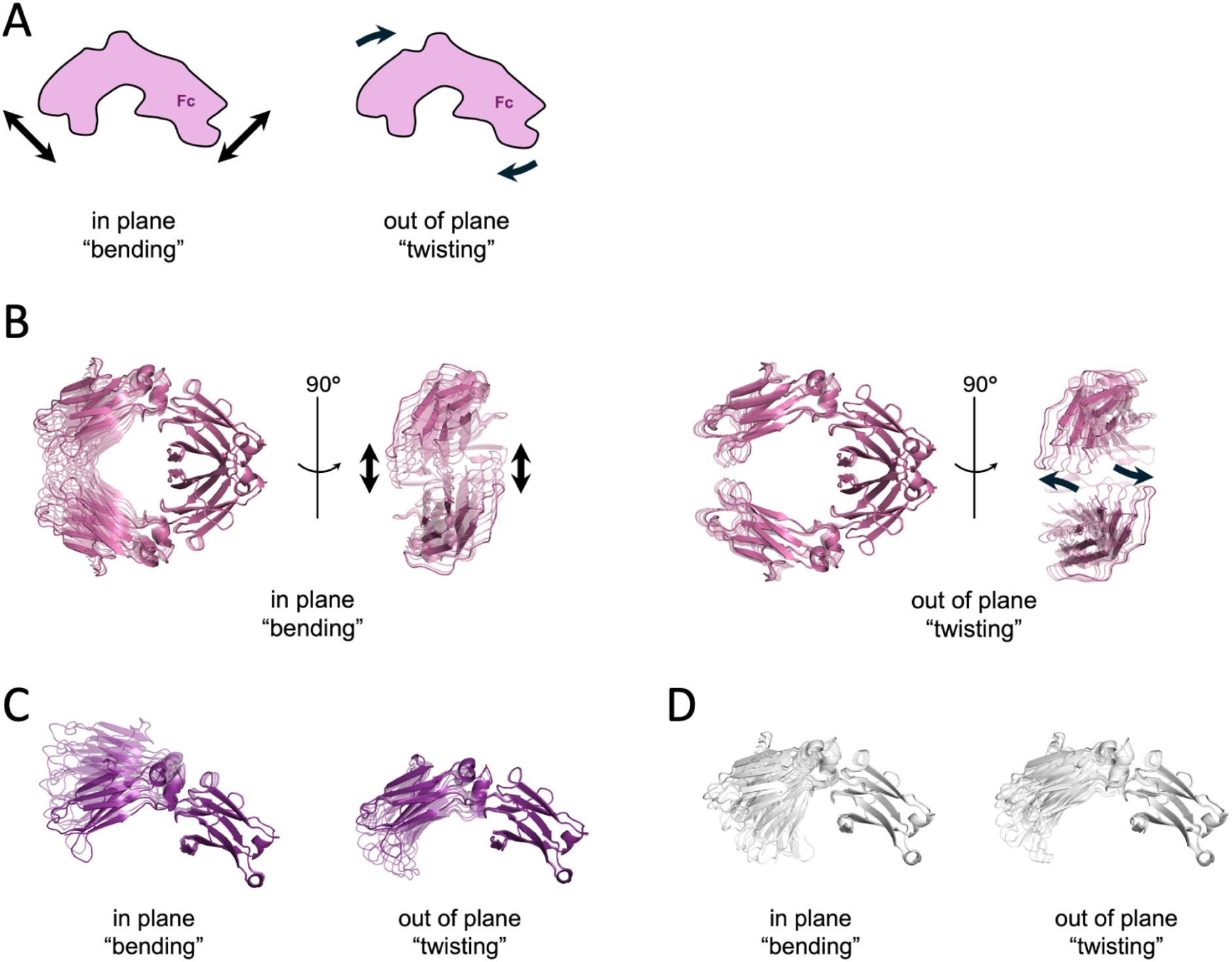
Normal mode analysis of the Fc antibody region. (A) Schematic representation of the lowest frequency mode (“bending”) and the second lowest frequency mode (“twisting”) of Fc. (B-D) Ribbon representations of normal mode motions in the Fc dimer structure (B, pink, PDBID: 4N0U), and two *wt* Fc monomers at low pH (C, purple, PDBID: 9CXL) and high pH (D, white, PDBID: 9CRT) calculated using NOLB (see Materials and Methods) and visualized via PyMOL. Structures are superimposed on the CH3 domain for easier visual comparison.

The three variant apo Fcs (M252H, M252R, and I253M) contained mutations at adjacent positions on the same structural loop of the CH2 domain. While the loop backbones in both low- and high-pH structures adopted conformations similar to the holo complex structure, we observed differences in sidechain orientations of the mutated residues attributable to shifts in relative domain orientation (Figure 3B). In the low-pH M252H and M252R Fc structures, the mutant His and Arg sidechains each interacted with negatively charged Fc residue E380 from the neighboring CH3 domain, forming a salt bridge and bringing the CH3 domain closer to the CH2 domain in these apo Fc structures compared to the holo conformation. Notably, this low-pH apo Fc conformation introduced clashes when superimposed onto the holo complex, between the Fc CH3 loop containing residue N434 and the H435 pH sensor, and the FcRn loop containing D130. However, these clashes were able to be resolved via loop relaxation (RMSD <0.2Å), and therefore would likely not preclude binding. Based on these observations, we concluded that at low pH, the salt bridge interaction between E380 and a positively charged Arg or His residue in place of a hydrophobic Met at Fc position 252 engenders a change in the apo Fc shape which must then adopt the holo conformation in order to bind FcRn efficiently.

To assess how these interactions behaved in solution, we performed 100 ns MD simulations for the M252H and M252R variants and quantified hydrogen bonds and salt-bridge formation involving E380, as presented in Figure 5. Both variants maintained direct contacts between residue 252 and E380 in roughly half of the trajectory frames. In addition, each variant showed frequent E380 interactions with K248, indicating that the negative charge of E380 is often stabilized by either residue 252, residue 248, or both simultaneously. These MD results reinforce the crystallographic findings by showing that protonated H252 or R252 engages E380 repeatedly in solution, stabilizing local conformations that alter the CH2-CH3 orientation and differ from those sampled by the *wt* Fc.

**Figure 5.**
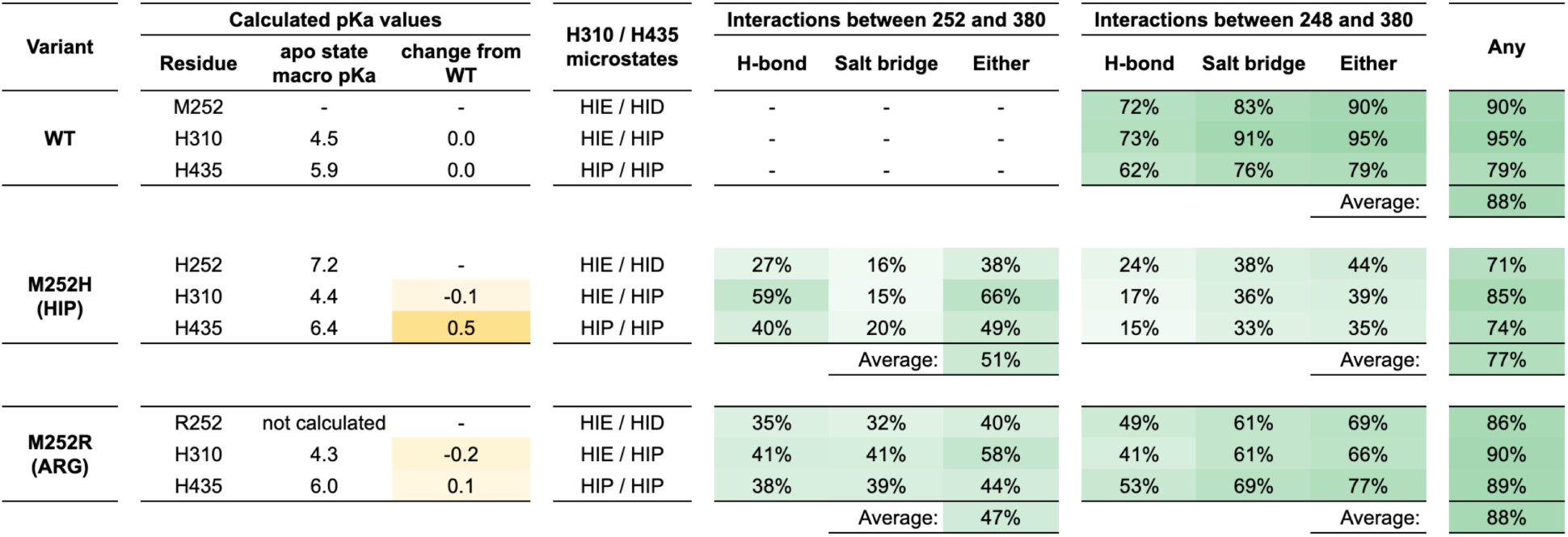
Calculated pKas and persistence of key interactions at the apo Fc CH2-CH3 domain interface in MD simulations of *wt*, M252H, and M252R variant apo Fc. FEP-calculated macro pKa values for His residues at positions 252, 310, and 435 are reported and used to determine 3 relevant microstates to simulate by MD. The fraction of MD frames exhibiting the indicated interactions between either positively charge HIP/ARG252 or LYS248 and GLU380 are shown for simulations set up with the indicated fixed protonation states.

The M252R, M252H, and I253M mutations in Fc weaken binding to FcRn, as indicated by a loss of 0.7-0.9 kcal/mol in binding free energy in SPR measurements. However, our initial standard-protocol FEP simulations, employing a thermodynamic cycle that assumed Fc to exist in the same conformational basin when bound to FcRn and in isolation, predicted these mutations to be strongly stabilizing. To account for the conformational variability seen in the crystal structures, we substituted solvent Δ*G* values calculated using an apo Fc model in place of the original, holo-conformation solvent Δ*G*. This change is illustrated in Figure 6, which compares the standard thermodynamic cycle (shaded region in the figure) with the modified version accounting for the conformational change, using the M252R mutation as an example. For mutations far from the CH2-CH3 interface, we expected the effect of this substitution to be minimal, at the noise level, but for key cases like M252R and M252H, we hypothesized that the effect could be substantial.

**Figure 6.**
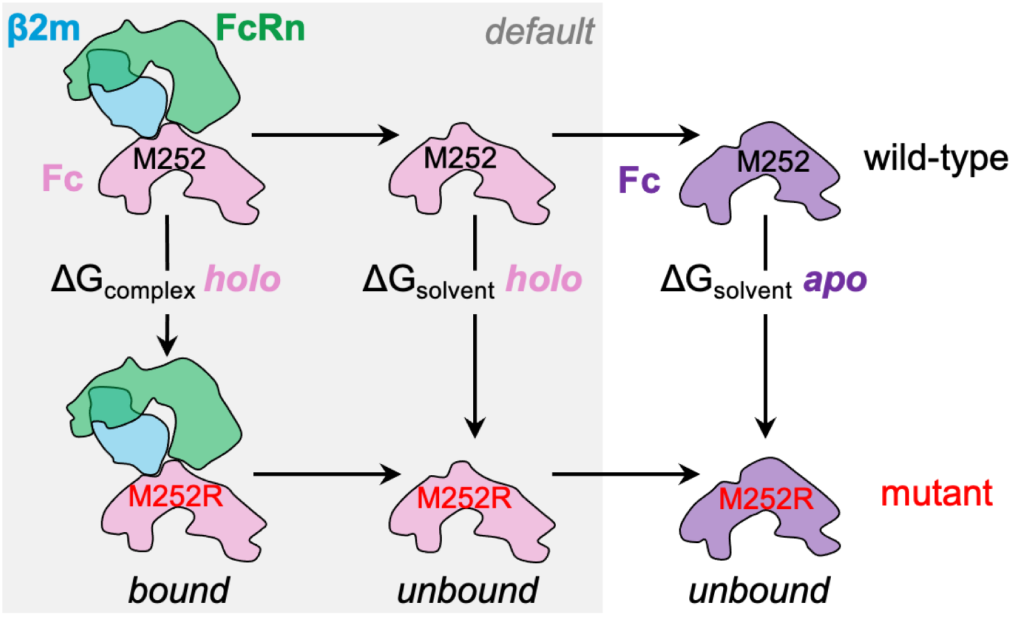
Thermodynamic cycle for FEP calculations of the M252R mutation in Fc protein and its effect on FcRn binding. Alchemical transformations for each mutation are simulated in two environments: the bound state, where Fc is in complex with the FcRn/β2m heterodimer; and the unbound state, where Fc is modeled alone. The default FEP protocol assumes that global Fc conformation is similar in both states. To address significant conformational changes observed in apo Fc crystal structures, we employed an extended thermodynamic cycle that explicitly accounts for structural differences.

That there was no clear relationship between pH and the conformational variation in the apo Fc structure (Supplemental Table 4), and the transitions among these structures aligned with the lowest-frequency, and therefore most easily accessible, normal modes (Figure 4) suggested that all of the observed apo and holo Fc conformations were likely to be sampled at room temperature, with comparable energies and similar population weights in the conformational ensemble. Therefore, we made the assumption that the transition between the apo and holo conformations for the *wt* protein incurs no energetic penalty. However, we noted the M252R and M252H mutations had the potential to shift the equilibrium toward the apo conformations, and it is these energy differences that we wished to uncover by using an apo-conformation starting model for Fc solvent leg simulations.

The low-pH apo Fc structures for the *wt*, M252H, and M252R variants were highly similar, with Cα RMSD below 0.4 Å. Therefore, we selected the highest-resolution of these crystal structures (2.2 Å, M252R low pH) to construct all-atom apo-conformation *wt* and YTE Fc models. We performed solvent leg-only FEP simulations for the full mutational dataset using these new apo Fc models and recalculated binding ΔΔ*G*s as the difference between the original (holo) complex leg Δ*G* and apo Fc solvent leg Δ*G*. For the M252H variant, in the dominant HIP protonation state (as determined using FEP-calculated macro pKa values of 6.3 and 8.3 in unbound and bound states, respectively, produced by the Protein FEP Groups output), the apo Fc solvent leg was ∼0.7 kcal/mol lower in energy than the corresponding holo Fc solvent leg, resulting in a less overbound ΔΔ*G* relative to experiment. The M252R variant exhibited a larger shift of ∼1.0 kcal/mol. With Groups treatment, the weighted average over the three possible His protonation states for M252H (HIP, HIE, HID) reduced the magnitude of the improvement compared to the single-state estimate, but overall, the use of the apo Fc model in the solvent legs reduced errors relative to experiment. After including the apo Fc solvent legs to account for conformational changes in Fc between bound and unbound contexts for all mutations, we observed that most cases were not affected by the change, many were improved to varying degrees, and relatively few cases produced somewhat larger errors, as indicated by the vertical tails shown in Supplemental Figure 3C.

Crystal structures of I253M variant Fc were solved at three different pH conditions (5.5, 6.5, 7.0), all of which were highly similar (RMSD <0.6Å), indicating no significant pH-dependent conformational changes. These structures also more closely resembled the *wt* structure at high pH rather than the *wt* at low pH (Supplemental Table 4). To further investigate the impact of apo conformational differences in computational simulations, we tested two different *wt* conformations in corrected solvent leg simulations: 1) the low-pH *wt* structure (as described above); and 2) the low-pH I253M structure. In each case, calculated solvent leg Δ*G* values remained very similar to the holo-conformation value (approximately -13.5 kcal/mol), which was expected given the largely unchanged local environment near the mutation across conformations. Because using the I253M-specific apo Fc model for the solvent leg did not improve the FEP prediction for this outlier case, in our final FEP results the reported ΔΔ*G* for the I253M Fc variant used the same *wt* apo Fc model as the rest of the dataset.

In the preceding paragraphs, we demonstrated improvement in predicted ΔΔ*G* values by accounting for global conformational differences in the unbound apo Fc structure relative to the holo complex. A second type of error due to incomplete sampling can arise if the FEP simulation is unable to access the lowest free energy mutant sidechain conformation. An issue of this type occurred for the T307W variant when using the low-pH apo crystal structure of the M252R mutation as a starting point, due to the presence of different rotamer conformations of the mutant Trp residue between apo and holo structures. This problem was readily remedied by modeling a different initial rotamer state for the sidechain, as discussed in Supplementary Results and accompanying Supplemental Figure 5, Supplemental Table 5, Supplemental Table 6, and Supplemental Table 7, and presented with the ΔΔ*G* correction shown as a vertical tail in Supplemental Figure 3D. We note that sampling errors of this sort could be avoided by incorporating state of the art conformational sampling algorithms when generating models for the mutant residue to locate the lowest mutant free energy structure, but as this procedure is rarely needed, these additional calculations are not part of the standard FEP workflow.

After incorporating this best-estimate binding ΔΔ*G* for the T307W variant, the results comprised the final retrospective FEP dataset presented in Figure 2 for all 88 experimental mutations. The *R*^2^ value of 0.38 and RMSE of 1.10 kcal/mol were comparable to the results obtained from previous protein binding FEP studies [17, 18, 27, 22]. Although a few predictions remained with absolute errors in the range of 2 kcal/mol, this could be expected in the distribution of FEP deviations from experiment for this size of dataset. The largest remaining outlier was I253M, for which we do not have an explanation—though for which we did remeasure the experimental *K*_D_, confirming the original value using different equipment and reagents (data not shown). Lastly, we note that the final outlier rate for the full Fc-FcRn dataset was less than 2%, which would not represent a barrier to practical antibody engineering applications.

## Discussion

Here we have presented a study of the Fc-FcRn system investigating the effects of amino acid mutations on binding affinity. We approached the system both experimentally through SPR affinity measurements and X-ray crystallography, and computationally via Protein FEP calculations and additional structural and trajectory analyses. The experimental binding affinity measurements presented here provide a substantial addition to the diversity of low-pH mutational binding data in the literature; in particular, the inclusion of several destabilizing variants extends the range of ΔΔ*G* values, allowing a more meaningful assessment of the correlation between predicted and experimental results. We expect the full mutational low-pH binding ΔΔ*G* dataset provided in the supplement to be useful in future studies of this important

By looking at specific cases that were improved at each stage of our calculations, we were able to identify important concepts to the function of the system. For example, the FEP-calculated pKa observed for Fc H435 in the bound context was higher using the OPLS5 force field compared to OPLS4, preferring HIP at pH 6.0. This is in better agreement with the consensus understanding of the mechanism for Fc-FcRn pH dependence, which depends on protonation of key Fc His residues H310 and H435 in the complex, and suggests that pi-pi or cation-pi interactions with nearby FcRn residue Trp131 may be critical to stabilizing the charged H435 sidechain. This is an important finding, and will undoubtedly be useful for other similar cases going forward.

Additionally, although not required for most cases, for certain mutations near or involving the key His residues, we found it was important to consider the effect of the mutations on the distribution of protonation microstates among these coupled titratable sites. In an extreme case, the half-life-extending DHS variant [14], which not only contains mutations near these two key histidines, but introduces two additional ionizable amino acid sidechains (D309 and H311) at the loop positions immediately before and after H310 in the primary amino acid sequence, was able to be successfully modeled—albeit at great computational expense—by considering all possible combinations of those protonation states. While this type of computation would not be recommended for most prospective cases in the current form, future development of FEP and adjacent free energy methods to improve the handling of such coupled sites has the potential to reduce computational cost and may be extremely useful in subsequent design efforts.

The inclusion of additional structural data and analyses allowed us to gain some insight into the dynamics of the system and how domain-level motions may play a role in binding and function. The experimental crystal structures of *wt* Fc and three Fc variants in the unbound apo state over a range of pH values, together with analysis of these new structures and previously reported Fc structures, demonstrated a degree of flexibility in the relative domain orientation of the antibody Fc region that appeared to be pH-independent. In contrast, local conformations appeared to depend both on pH and on the specific amino acid sequence of the variant, suggesting that adoption of the holo conformation capable of binding to FcRn may also favor (or be favored by) certain mutations.

While the standard FEP protocol uses the same input structure for both the solvent and complex legs of the calculation, we found that using an apo Fc model specific to the unbound state yielded substantial improvement for a number of cases. However, not all cases improved, which may reflect differences in preferred global conformations for different mutations; we note that when using a single apo model for all mutations that we also assumed that the reorganization energy to adopt this conformation from the binding-competent conformation present in the complex structure is the same for all variants, which may not be the case in the physical system. With this in mind, we propose that a more complete understanding of the relative motion of Fc domains, in particular via the “breathing” or “twisting” normal modes, may be useful in Fc engineering and warrants further study.

With regard to the remaining outlier cases, we note that Fc variants M252H, M252R, and I253M are mutations of the loop containing the YTE mutations in the original crystal structure. Although we could not identify any obvious deficiencies in the all-atom *wt* model, it is possible that some subtle difference between the YTE crystal structure and the true *wt* conformation, inherited by our all-atom models, prevented sampling of a conformation relevant to the *wt* complex. Additionally, the only two other cases with errors larger than 2 kcal/mol were the Fc V259I/V308F/M428L (IFL) triple mutant, and the YTE Fc S254V variant. The IFL variant contained the M428L mutation from the LS variant, which is structurally near the location of the YTE Tyr252 sidechain, and S254V is also located on this particular loop. Further structural investigations of a human Fc-FcRn complex with a *wt* amino acid sequence for this loop may prove useful in determining whether these outliers may be due to structural differences in this loop region among the different variants in complex with FcRn.

We adopted an iterative approach to improve the agreement of computational predictions with experimental results, identifying potential deficiencies in our models and simulation protocols, and updating them based on new experimental and computational results. This methodology, similar to the typical Design-Make-Test-Analyze (DMTA) cycle used in drug discovery, may be just as readily applied in a prospective project as it is in the current retrospective study, with the understanding that a fraction of initial FEP predictions may exhibit significant errors due to incomplete sampling. The present work demonstrated the successful use of this iterative method by addressing outlier cases for which predicted ΔΔ*G* values deviated from experimental result, through further experimental studies and refinement of structural models and simulation parameters.

In summary, despite the challenges associated with predicting binding free energies at acidic pH for a system with a high density of titratable residues at the binding interface, including two histidines that are expected to undergo a change in protonation state as part of the pH-dependence mechanism, our FEP-based study was able to accurately reproduce experimental binding ΔΔ*G* values for a large dataset of single, double and triple mutant Fc-FcRn variants. By explicitly addressing the effects of coupled mutations on nearby titratable sites and accounting for conformational flexibility in unbound Fc, we were able to achieve a level of accuracy that has come to be expected from FEP results, and which is necessary for prospective work, thus opening the door for further efforts toward rational, *in silico* design of antibody Fcs with increased half-life.

## Materials and Methods

### Preparation of recombinant IgG, FcRn, and gp120 core for binding experiments

Expression vectors for Fc variants selected for experimental K_D_ measurement were generated by site-directed mutagenesis (GeneImmune) using the heavy chain expression vector for antibody VRC01 as a template [26]. Expression vectors for the recombinant soluble human FcRn α chain, the β2m chain with a C-terminal 8x His-tag, and clade A/E 93TH057 HIV-1 gp120 core(e) were constructed as described previously [28, 29]. Wild-type and variant Fc were expressed as IgG by transiently transfecting Expi293F cells (Thermo Fisher Scientific) with equal amounts of heavy and light chain expression plasmids using Turbo293 transfection reagent (SPEED Biosystem) according to manufacturers’ protocols. Wild-type and variant FcRn and gp120 core(e) were also produced in Expi293F cells by transient transfection. At 5 days post-transfection, cell culture supernatants were harvested and purified using Protein A Sepharose CL-4B resin (GE Healthcare) for Fc variants, cOmplete His-Tag purification resin (Sigma Aldrich) for FcRn and its variants, and 17b antibody-conjugated affinity column chromatography for gp120 core(e). The proteins were further purified by size-exclusion chromatography (SEC) using a HiLoad 16/600 Superdex 200 prep grade column (GE Healthcare).

### K_D_ measurement for IgG-FcRn binding by surface plasmon resonance

For SPR measurements, a CM5 sensor chip was coated with clade A/E 93TH057 HIV-1 gp120 core(e) immobilized by amine coupling and used to capture *wt* or Fc-variant VRC01 IgG to ∼80-130 resonance units (RU) using Biacore T200 (GE Healthcare). Recombinant soluble *wt* or variant FcRn analytes were injected into the reference cell and the sample flow cell with the immobilized-IgG surface at concentrations ranging from 7.8 to 4000 nM or 7.8 to 10,000 nM in HBS-EP+ buffer (GE Healthcare) supplemented with 20 mM sodium citrate, pH 6.0, at a flow rate of 50 μL/min. Association and dissociation phases were monitored for 2 and 1 min, respectively. All SPR experiments were performed at 25°C. Reference-subtracted curves were fitted to a 1:1 Langmuir binding model or steady-state affinity model using Biacore T200 Evaluation Software 3.1.

### Calculation of ΔΔG from K_D_ values

We converted binding affinities (*K*_D_) to binding free energy (Δ*G*) values for each mutant using the standard relationship between dissociation constant and binding free energy, Δ𝐺 = −𝑅𝑇 ln(𝐾_D_/𝐶), where R is the ideal gas constant (kcal mol^-1^ K^-1^), T is temperature (K), and C is the standard reference concentration of 1 M. Binding ΔΔ*G* values were computed for the variants in each publication by subtracting the *wt* Δ*G* from the Δ*G* of each corresponding mutant within the same study. For publications reporting measurements of the same Fc variant in different antibody (i.e. Fab) contexts, we reported a different ΔΔ*G* value for each unique variant-antibody combination, relative to the *wt* Fc measurement for the same antibody. ΔΔ*G* values of variants with measurements reported in multiple publications were averaged in the final dataset.

### Protein expression and purification for Fc crystallization

DNA sequences encoding *wt* IgG1 Fc, and M252H, M252R, and I253M Fc variant proteins were cloned into the mammalian expression vector pVRC8400. Point mutations were introduced into the constructs using KOD Hot Start polymerase (Novagen). The constructs were expressed in Expi293™ cells (Invitrogen), with polyethylenimine (Polysciences) used as the transfection reagent. Growth media were collected five days post-transfection, and secreted Fc proteins were purified using Protein A affinity chromatography, followed by SEC in PBS buffer at pH 7.4.

### Determination of apo Fc crystal structures

SEC fractions containing purified Fc proteins were pooled and concentrated to 10.5 mg/mL (*wt* Fc), 12.5 mg/mL (M252H Fc), 15.0 mg/mL (M252R Fc), or 9.0 mg/mL (I253M) in the SEC buffer. Screening for initial crystallization conditions was carried out in 96-well sitting drop plates using the vapor-diffusion method after setup with a Mosquito crystallization robot (TTP LabTech) using various commercially available crystallization screens: JSCG+ (Qiagen), MSCG-1 (Anthracene) and LMB and Proplex HT (Molecular Dimensions).

Diffraction quality crystals were obtained after 2 days under multiple conditions. For each variant, crystals from 2 or 3 crystallization conditions were selected for data collection, specifically including crystals that were grown at neutral/high pH or at acidic pH. For *wt* Fc: 0.1 M sodium citrate pH 5.5 (“WT low pH” structure, PDBID: 9CXL); and 0.1 M HEPES pH 7.0, 20% PEG 20,000, and 1 M ammonium sulfate (“WT high pH,” 9O7A). For M252H Fc: 0.1 M sodium citrate pH 5.6, 20% propanol, and 20% PEG 4000 (“M252H low pH,” 9D09); and 0.1 M Bis-tris propane pH 7.5 and 8% PEG8K (“M252H high pH,” 9CY6). For M252R Fc: 0.1 M sodium citrate pH 5.6, 20% propanol, and 20% PEG 4000 (“M252R low pH,” 9D06); and 0.1 M Tris pH 8.0, 0.08 M sodium formate, and 7.5% PEG 20,000 (“M252R high pH,” 9D9Q). For I253M Fc: 0.1 M citrate pH 5.5 and 15% PEG 6000 (“I253M pH 5.5,” 9O64); 0.1 M MES pH 6.5, 10% PEG 5000 and 12% propanol (“I253M pH 6.5,” 9O75); and 0.1 M MES pH 7.0, 15% PEG 20,000 and 12% propanol (“I253M pH 7.0,” PDBID: 9O78). Prior to data collection, crystals were cryoprotected in mother liquor supplemented with 40% ethylene glycol and flash frozen in liquid nitrogen.

X-ray diffraction data were collected at 100 K on beam line 17ID-1 (AMX) or 17 ID-2 (FMX) at the National Synchrotron Light Source II at Brookhaven National Lab (NY). Diffraction data were processed with autoPROC and scaled using AIMLESS from the CCP4 software suite [30, 31, 32]. Molecular replacement was performed with PHASER [33], using the Fc chain from PDB 6WNA as a search model [23]. Manual rebuilding of the structures using COOT [34] was alternated with refinement using Phenix.refine [35]. The MolProbity server [36] was used for structure validation and PyMOL was used for structure visualization and RMSD analysis [37].

### All-atom model preparation from crystal structures

Initial structural coordinates were obtained either from previously reported (YTE Fc-FcRn complex, PDB 6WNA) [23] or newly solved structures (apo Fc variants). For the complex structure, limited structural modeling and crystallographic refinement were performed using COOT [34] and Phenix [35], including the addition of ordered solvent near the binding interface. Bond orders were assigned and hydrogens added using the Protein Preparation Wizard panel (PPW) in Maestro (Schrödinger). Initial pKas were calculated and protonation states assigned within the PPW using PROPKA at pH 6.0 [38]. The YTE Fc chain from the 6WNA model was IgG4 subtype, and was monomeric due to 6 mutations disrupting the CH3-CH3 dimerization interface [23]. We mutated 17 residues in total to convert the model to the YTE IgG1 amino acid sequence. The required covalent bonds were modeled between sugar monomers, and any water molecules beyond 3 Å from the nearest protein atom were removed. Finally, a model of the *wt* Fc-FcRn complex was created from the YTE complex model by simple mutation of residues Y252, T254, and E256 in the YTE Fc chain to their *wt* counterparts, namely M252, S254, and T256, using the Maestro GUI. A more detailed accounting of all-atom model preparation is given in Supplementary Methods.

### FEP-based pKa calculation and pH-specific protonation state assignment

Protein FEP calculations were run using OPLS4 or OPLS5 to calculate titratable residue pKas as described previously [19]. Output perturbation map (.fmp) files and experimental pH (6.0) were used to determine pH-specific populations of each protonation microstate, variant Δ*G* values, and residue pKas following the Protein FEP Groups approach [22] via the protein_fep_groups.py script in the Schrödinger Suite. Protonation states in the input complex models were updated accordingly to create force field- and pH-specific models, with protonation states with the highest population in the bound complex state selected for each site.

### Retrospective FEP calculations

Protein FEP calculations were run for each of three data subsets (mutations of *wt* Fc, YTE Fc, or *wt* FcRn) with simulation times of 10 ns and OPLS5 force field using default protocols from the Schrödinger Suite 2024-2 release. Multiple mutations were performed sequentially according to the perturbation map automatically generated by the Protein FEP workflow. All alternate protonation states were included for perturbations to or from ionizable residues in the FEP map, and the Protein FEP Groups treatment was applied to calculate pH-specific predicted Δ*G* values for each variant at experimental pH 6.0. Although several measurements in the experimental dataset were actually reported using pH 5.8 as shown in Supplemental Figure 6, because FEP-predicted Δ*G* values after Protein FEP Groups treatment at pH 5.8 for those cases differed only negligibly from the pH 6.0 predicted Δ*G* values (by <0.1 kcal/mol in each case), only pH 6.0 predicted values are reported here. Likewise, the pH 5.8 experimental Δ*G* values were included as-is alongside corresponding pH 6.0 measurements when calculating average per-variant experimental Δ*G*s.

### Selection of variants for analysis of coupling to pH-sensor His residues

Putative pH-sensing Fc histidine residues H310 and H435 were selected for analysis to determine how their bound or unbound pKas and effect on overall predicted binding ΔΔ*G* were influenced by nearby mutations. Variants with potential coupled sites were identified for additional FEP calculations based on the following protocol. From the mutations.txt file used to prepare each FEP map (*wt* Fc, YTE Fc), “neighbor mutation sites” were selected which contained any sidechain atom within a given cutoff distance of any sidechain N or O atom in the coupling sites in the input all-atom model (in this case the relevant atoms were the H310 and H435 imidazole nitrogens). A cutoff of 6.0 Å was used for mutations where the neighboring residue was mutated to or from a charged or ionizable residue (D/E/H/K/R); a closer cutoff of 4 Å was used for neutral mutations. Variants containing any mutation at a selected site were included in separate FEP perturbation maps with all combinations of H310 and H435 protonation microstates, as well as the protonation microstates of ionizable residues at mutated sites (either *wt* or mutant), as described in Supplementary Methods.

### Calculation of solvent leg ΔG values using new crystal structure models

All-atom models of *wt* and YTE apo Fc were prepared starting from the crystal structure coordinates of the M252R low pH structure (PDB ID: 9D06). Residues 252, 254, and 256 in the Fc chain were mutated to either the *wt* (Met, Ser, Thr) or YTE (Tyr, Thr, Glu) sequences, respectively, to serve as input for subsequent apo-state FEP calculations. After preparation using the Maestro PPW as above, protonation states of all titratable residues were adjusted to match those in the pH 6.0 holo *wt* Fc model.

Solvent leg calculations for the retrospective set of mutations were performed using the *wt* or YTE apo Fc model using the same parameters used for the holo complex, except the calculation was run using the Protein FEP thermostability (rather than protein binding) protocol, as the FcRn binding partner was not present. The resulting solvent Δ*G* values were used to compute modified binding ΔΔ*G* values via the formula: 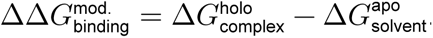.

### Normal mode analysis of Fc structures

Low frequency normal modes, describing collective atomic motions typically associated with the most flexible regions, were calculated for all monomeric holo and apo Fc structures as well as for a dimeric YTE Fc structure (prepared from PDB 4N0U via crystallographic symmetry) using a computationally efficient NOn-Linear rigid Block normal mode analysis approach (NOLB) [39]. Normal modes for individual structures were computed using the command `./NOLB structure.pdb`, and transitions between two structures using the two lowest-frequency modes were generated with the command `./NOLB structure1.pdb structure2.pdb -n 2 -s 20`.

### Data analysis and figure and manuscript preparation

Data analysis was done using Python [40] and R [41]. Plots were generated in R using the ggplot2 package [42]. Structural figures were prepared using PyMOL [37]. The manuscript was prepared using Quarto and RStudio [43, 44].

## Data availability

Experimental data, prepared all-atom structural models, and FEP results are available in the GitHub repository accompanying this manuscript, accessible via Zenodo [45] at DOI: 10.5281/zenodo.19777392.

## CRediT authorship contribution statement

**Jared M. Sampson:** Conceptualization, Data Curation, Formal analysis, Investigation, Methodology, Software, Visualization, Writing - Editing & Review, Writing - Original Draft. **Alina P. Sergeeva:** Data Curation, Formal analysis, Investigation, Methodology, Visualization, Writing - Editing & Review, Writing - Original Draft. **Tianyang Gao:** Data Curation, Formal analysis, Investigation, Methodology, Visualization, Writing - Editing & Review, Writing - Original Draft. **Young Do Kwon:** Investigation, Visualization, Writing - Editing & Review. **Eswar Reddem:** Investigation, Visualization, Writing - Editing & Review. **Fabiana A. Bahna:** Investigation, Writing - Editing & Review. **Seetha M. Mannepalli:** Investigation, Writing - Editing & Review. **Baoshan Zhang:** Investigation, Writing - Editing & Review. **Peter D. Kwong:** Funding acquisition, Resources, Supervision, Writing - Editing & Review. **Lawrence Shapiro:** Conceptualization, Funding acquisition, Resources, Supervision, Writing - Editing & Review. **Barry Honig:** Conceptualization, Funding acquisition, Methodology, Resources, Supervision, Writing - Editing & Review. **Richard A. Friesner:** Conceptualization, Funding acquisition, Investigation, Methodology, Resources, Supervision, Writing - Editing & Review.

## Acknowledgements

The work was supported by the Gates Foundation, INV-080104 (B.H., L.S., and R.A.F.), by the NIH grant R35-GM1395858 (B.H.), and by the Intramural Research Program of the Vaccine Research Center, National Institute of Allergy and Infectious Diseases, National Institutes of Health.

This work was supported in part by the Gates Foundation. The conclusions and opinions expressed in this work are those of the author(s) alone and shall not be attributed to the Foundation. Under the grant conditions of the Foundation, a Creative Commons Attribution 4.0 License has already been assigned to the Author Accepted Manuscript version that might arise from this submission. Please note works submitted as a preprint have not undergone a peer review process.

This research was supported in part by the Intramural Research Program of the National Institutes of Health (NIH). The contributions of the NIH author(s) are considered Works of the United States Government. The findings and conclusions presented in this paper are those of the author(s) and do not necessarily reflect the views of the NIH or the U.S. Department of Health and Human Services.

## Supplementary Information

### Supplementary Methods

#### All-atom Fc-FcRn structural model preparation and protonation state assignment

We selected the X-ray crystal structure of a monomeric YTE variant human IgG4 Fc chain bound to the human FcRn+β2M heterodimer (PDB 6WNA, 2.4 Å resolution) as the starting point for our study [23]. We performed limited structural modeling and crystallographic refinement including the addition of visible ordered water molecules near titratable residues near the binding interface and an increase in the percentage of Ramachandran-favored residues from 95.48% to 98.58%. The final crystallographic model gave improved overall refinement statistics, reducing *R*_work_/*R*_free_ from 0.214/0.263 in the original PDB entry to 0.206/0.239 in the re-refined version.

We introduced 17 point mutations into the re-refined model in Maestro (Schrödinger, Inc.) to convert the Fc chain from IgG4 to the IgG1 subtype used for the experimental binding data. These included reversion of 6 mutations at the CH3-CH3 interface that had been introduced by Shan and coworkers to produce monomeric Fc [23]. Notably, none of these mutations involved residues at the FcRn binding interface. Finally, from this IgG1 YTE variant Fc-FcRn model, we made three mutations (Y252M/T254S/E256T) to revert the YTE residues to the *wt* sequence. The resulting united-atom *wt* and YTE variant IgG1 Fc-FcRn models were used as the basis for the preparation of all-atom models for use in FEP calculations.

Using the Protein Preparation Wizard in Maestro, we added hydrogens, assigned bond orders, and configured initial protonation states at pH 6.0 with PROPKA3 [38]. The resulting preliminary all-atom *wt* and YTE models were used as inputs to Protein FEP to calculate pKas for key interface titratable residues using either the OPLS4 or OPLS5 force fields, perturbing each interface titratable site from the initial modeled state to its alternate protonation states (e.g. H:HIP310 to HID and HIE). The pKa values and the accompanying microstate populations reported after Protein FEP Groups treatment at pH 6.0 were used to determine pH-specific protonation states for each model according to which state was predicted to be dominant in the bound complex. Interestingly, FEP-based pKa calculations using the fixed-charge OPLS4 force field predicted Fc residue H435 to be in the neutral HID tautomer in the bound state; however, the OPLS5 calculations for both *wt* and YTE models each yielded higher pKa values for H435, indicating the positively charged HIP microstate was preferred. As a final step, the indicated force field-specific protonation states were then incorporated into the models, resulting in the fully prepared pH 6.0 *wt* and YTE models ready for use in retrospective FEP calculations.

#### Accounting for coupled titratable sites in FEP via explicit multiple mutations

The Protein FEP Groups treatment was previously described in ref. [22] from the main text, and is based on a similar approach from Oliveira et al. [24]. For completeness, a brief summary of the methodology is included here, along with a description of how it can be used to treat coupling among titratable sites.

Protein FEP+ calculations involve alchemical transitions between fixed-topology models, so no protonation or deprotonation occurs over the course of the simulation in a time-dependent manner. However, by including FEP+ edges in the perturbation map in a way that provides alchemical perturbation paths from the microstate present in the starting model to all possible relevant protonation microstates for a given ionizable site or set of sites, from a single FEP+ map we can determine relative ΔΔ*G*s among all of the microstates, and therefore fractional populations of each microstate (as represented by a node in the FEP perturbation connectivity graph) in the unbound and bound physical macrostates at a given pH.

To account for protonation state effects on binding robustly, each perturbation requires three separate simulations, or “legs,” representing three physical macrostate contexts: bound state (complex leg); unbound state (solvent leg) and model state (fragment leg, run as a single, capped amino acid residue). From these simulations we obtain predicted Δ*G* values relative to the input structure for each microstate in each context. Using a single amino acid for the model leg allows the use of known pKas from the literature for the amino acids in this reference state to calculate model populations for each titratable site being considered using the Henderson-Hasselbalch equation (Equation 2):

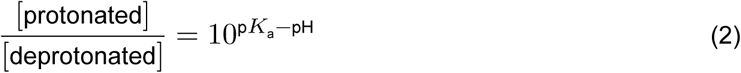

Given these literature-derived starting populations, if we consider the model, solvent, and complex as states 0, 1, and 2, respectively, the fractional solvent population (𝑝*_i_*^1^) for each microstate 𝑖 can be calculated as a function of the model population (𝑝*_i_*^0^) and the differences among the predicted folding free energies 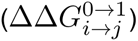 for all nodes 𝑗 (including 𝑖) in the same node group using Equation 3:

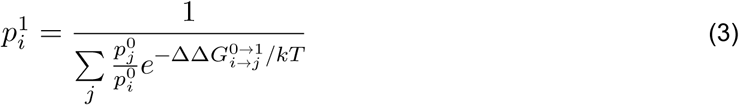

where 𝑘 is the Boltzmann constant and 𝑇 is the temperature in K. Similarly, the complex population for the node (𝑝*_i_*^2^) can then be calculated from the solvent population using the same equation with the solvent populations and binding free energies as input (i.e. increment the indices for all superscripts).

With solvent (𝑝*_i_*^1^) and complex (𝑝*_i_*^2^) populations in hand, the overall pH-specific binding free energy of a given variant 𝑋 can be calculated by applying appropriate state penalties based on the fractional populations of the starting and ending states via:

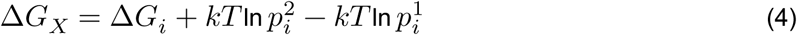

For complex scenarios where multiple titratable sites must be considered, we treat the sites in one of two ways. First, if a titratable residue in the starting model is mutated to another amino acid in the perturbation map, it is perturbed to its alternate states in the WT node group, but this perturbation is not repeated in other variant node groups. In this case, the unbound and bound populations of that titratable residue in the *wt* context are applied as a factor to the unbound and bound populations of all variant node groups which retain that site. That is, the effects of titratable sites from the WT node group are inherited by the other variants if present, and assumed not to be affected by mutations in the variant. Furthermore, different sites in the WT node group are also assumed to be independent of one another, as a full combinatorial accounting of the microstates resulting from all titratable sites would be computationally intractable.

For specific cases, we employ a second approach, where coupling between titratable and/or mutation sites is explicitly defined by specifying a multiple mutation in the FEP+ input mutations file. For example, the mutation N434Y coupled to the pKa of *wt* titratable residue H435 would require having predicted Δ*G* values for the N434Y mutation in the context of each of the 3 His states at position 435. In the input mutations.txt file, with the HIP435 protonation state in the starting model, this could be written as the following:

`H:434->TYR`

`H:435->HID`

`H:435->HIE`

`H:434->TYR,H:435->HID`

`H:434->TYR,H:435->HIE`

This input would result in 6 nodes total:

`WT`

`H-HIP435HID`

`H-HIP435HIE`

`H-ASN434TYR`

H-ASN434TYR,H-HIP435HID

H-ASN434TYR,H-HIP435HIE

with the first 3 nodes comprising the WT node group, and latter 3 nodes the H-N434Y node group. (Note that in the input mutations format, mutations separated by commas are automatically expanded by the FEP+ workflow to include all combinations of the component mutations, so although instructive here, the first 3 lines of the mutations.txt above are optional, as they are implied by the double-mutation lines.)

This process may also be applied to coupled titratable sites. For example, considering the interplay between the M252H mutation and *wt* Fc residues H310 and H435, both modeled as HIP, the mutations file used in our simulation included the following:

H:252->HID,H:310->HID,H:435->HID

H:252->HID,H:310->HID,H:435->HIE

H:252->HID,H:310->HIE,H:435->HID

`H:252->HID,H:310->HIE,H:435->HI`

H:252->HIE,H:310->HID,H:435->HID

H:252->HIE,H:310->HID,H:435->HIE

H:252->HIE,H:310->HIE,H:435->HID

H:252->HIE,H:310->HIE,H:435->HIE

H:252->HIP,H:310->HID,H:435->HID

H:252->HIP,H:310->HID,H:435->HIE

H:252->HIP,H:310->HIE,H:435->HID

H:252->HIP,H:310->HIE,H:435->HIE

It should be noted that the simplifying assumption, that the perturbations sampled in FEP do not affect (and are not affected by) the protonation state of any other residue in the structure, is successful most of the time because the results produced by protein mutation FEP are *relative* free energy changes. Even if a protonation state (or change thereof) is important to binding, having an incorrect protonation state modeled will, in general, have minimal effect on most results because the calculations benefit from cancellation of error when both the *wt* and mutant simulations suffer from the same modeling error and the output result is reasonable. However, if the pKa of a nearby residue is tightly coupled to a perturbation (either a titration between microstates of a residue, or a mutation to a different amino acid), the errors are no longer balanced and the effect on predicted Δ*G* values can be substantial, as seen, for example, with the DHS variant discussed in the main text.

### Supplementary Results

#### Correction of the T307W outlier arising from convergence challenges

The T307W mutation was moderately unfavorable according to SPR measurements (binding ΔΔ*G* = 0.8 kcal/mol). Using the default FEP protocol—where the holo Fc conformation was employed in the solvent leg (see Figure 6)—we predicted a binding affinity change of -0.8 kcal/mol, resulting in an error of 1.6 kcal/mol. Notably, using the apo conformation in the solvent leg further increased the prediction error by approximately 1 kcal/mol. This result was unexpected, as the T307 residue is located in the CH2 domain and does not interact with the CH3 domain in either the holo or apo conformations (see Supplemental Figure 5A). Therefore, the choice of solvent leg conformation was not expected to significantly impact the prediction for this mutation, let alone exacerbate the error.

To investigate the anomalous behavior of the T307W mutation, we conducted three independent simulations using either the default or “extended” FEP protocol, that is, using either holo or apo conformation starting models in the solvent leg, respectively. Solvent leg energies exhibited substantial fluctuations in both cases, ranging from -4.76 to -6.33 kcal/mol for the holo conformation and from -3.97 to -4.77 kcal/mol for the apo conformation, indicating poor convergence across runs.

Since solvent leg energy was sensitive to the conformational states sampled, we examined pairwise interactions between T307W and nearby residues (within 6 Å) across all runs (Supplemental Table 5). The lowest solvent leg energy (-6.33 kcal/mol) was associated with stabilizing interactions: a hydrophobic contact between the methyl group of T256 and the indole ring of T307W (observed in 98% of the trajectory), a cation-pi interaction between K288 and the aromatic sidechain of T307W (98%), and a hydrogen bond between E258 and the N-H group of the T307W indole ring (65%). A representative structure is shown in the right panel of Supplemental Figure 5B. These sidechain interactions strongly correlated with more favorable solvent leg energies (Supplemental Table 5).

The local environment of the mutation site differed between the holo and apo structures (Supplemental Figure 5B). Residue T307 adopted distinct rotameric states in each, corresponding to the two most frequent rotamers found in crystal structures (46.8% in apo, 41.8% in holo). T256 also differed in orientation: in holo, it promoted a hydrophobic interaction with T307; in apo, its hydroxyl group instead faced T307, disrupting this contact. These differences contributed to less favorable solvent leg energies in the apo conformation, as reflected in the reduced frequency of T256-T307W interactions (Supplemental Table 5).

To confirm that local environment differences—and not interdomain angle—drove the increased error with the apo structure, we ran reverse FEP simulations (Trp307→Thr) starting from mutant structures pre-stabilized in their local environment (“holo stab MT” and “apo stab MT”; Supplemental Figure 5C). Both simulations adequately sampled the stabilizing state and resulted in near-degenerate solvent leg energies, 5.38 and 5.61 kcal/mol, supporting the conclusion that the local environment was responsible for the discrepancy.

Next, we investigated whether one could improve sampling of the stabilizing state in the mutant trajectories by varying the rotamer state of the mutated residue. Three starting geometries were constructed for both holo and apo conformations, using the three most common Thr rotamers for T307. After manually selecting each rotamer, we minimized the mutated residue and its neighbors within 6 Å. Forward simulations of T307→W (Supplemental Table 6) revealed that only rotamer2 in holo conformation led to sampling of the stablizing state, yielding a solvent leg energy of -5.25 kcal/mol. Similarly, reverse simulations using various Trp rotamers yielded stabilizing interactions only in the apo rotamer2 case, with a corresponding energy of -5.46 kcal/mol. The range of solvent leg Δ*G* across rotamers 1-3 exceeded 1 kcal/mol for the T307→W mutation (Supplemental Table 6), likely due to the residue size change upon mutation. In contrast, mutations to residues closer in size to Thr (e.g. T307→Q,E) showed smaller solvent leg energy variations (0.5-0.6 kcal/mol), consistent with the stochastic range of FEP. Thus, undersampling was more pronounced for small-to-large mutations like T307W, and varying rotamers alone may be insufficient. One potentially more robust approach may involve sampling multiple rotameric starting points and selecting the mutant structure yielding the most favorable energy, assuming convergence in the *wt* runs.

Other mutations involving large sidechain size increases in our dataset—such as N434W and M252W—exhibited only modest solvent leg energy variation (0.2-0.3 kcal/mol) between holo and apo conformations Supplemental Table 7, suggesting the larger error seen for T307W was due to local structural sensitivity.

In this poorly converging case, we enhanced sampling through repeated simulations and identification of the most stable mutant state. Using the lowest solvent leg energies from three independent runs, we computed ΔΔ*G* of the T307W mutant as Δ*G*complex - Δ*G*solvent = -6.01- (-6.33) = 0.32 kcal/mol, which reduces the prediction error from 1.6 to 0.5 kcal/mol, improving agreement with experiment (ΔΔ*G* = 0.8 kcal/mol).

### Supplementary Tables

**Supplemental Table 1.**
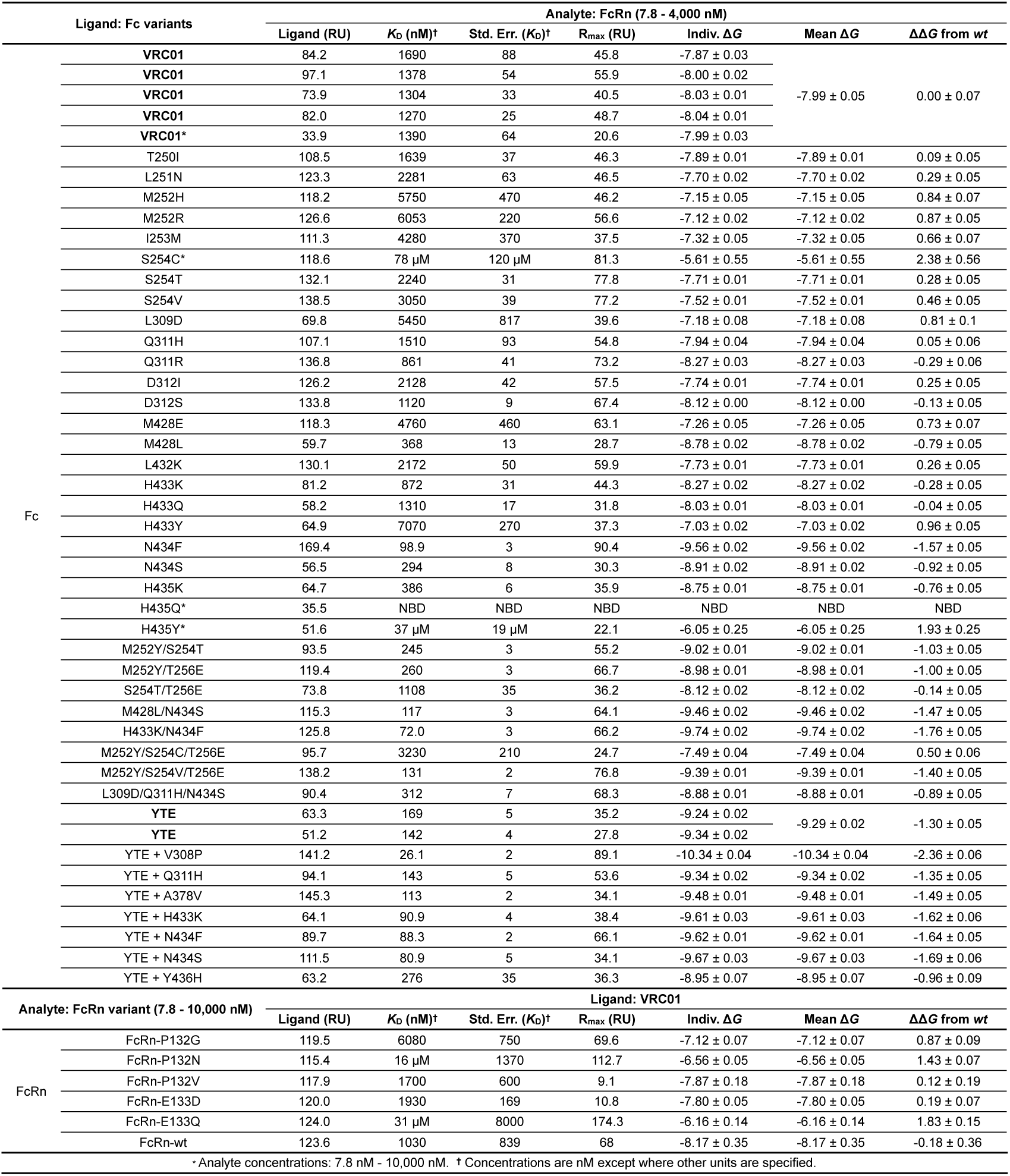
Equilibrium *K*_D_ and binding ΔΔ*G* values for *wt* and variant Fc-FcRn complexes at pH 6.0. Values for the binding of soluble FcRn analyte to immobilized VRC01 IgG from the measurements in Supplemental Figure 1. *K*_D_ values and standard errors based on fits of steady-state responses or kinetic parameters over the indicated concentration series are indicated. Immobilized IgG ligand densities and estimated maximum response (R_max_) are specified in resonance units (RU). Binding Δ*G*s are calculated for each measurement and averaged per variant, and resulting ΔΔ*G*s from the mean *wt* value are indicated.

**Supplemental Table 2.** Observed changes in predicted Binding ΔΔ*G* values (kcal/mol) for 7 outlier cases before and after the inclusion of explicit coupling to Fc H310/H435 protonation states in FEP calculations.

| Variant | Expt. $\Delta\Delta G$ | Orig. Pred. $\Delta\Delta G$ | Coupled Pred. $\Delta\Delta G$ | $\Delta$ Abs. Err. |
| --- | --- | --- | --- | --- |
| L251N | 0.32 | 0.62 | 0.64 | 0.02 |
| M252H | 0.87 | -1.40 | -1.95 | 0.55 |
| M252R | 0.90 | -2.08 | -1.92 | -0.16 |
| I253M | 0.69 | -2.43 | -1.79 | -0.64 |
| H435K | -0.73 | -2.50 | -1.79 | -0.71 |
| H435Y | 1.96 | 1.14 | 1.23 | -0.09 |
| L309D,Q311H,N434S | -0.91 | 2.78 | 1.01 | -1.77 |

**Supplemental Table 3.**
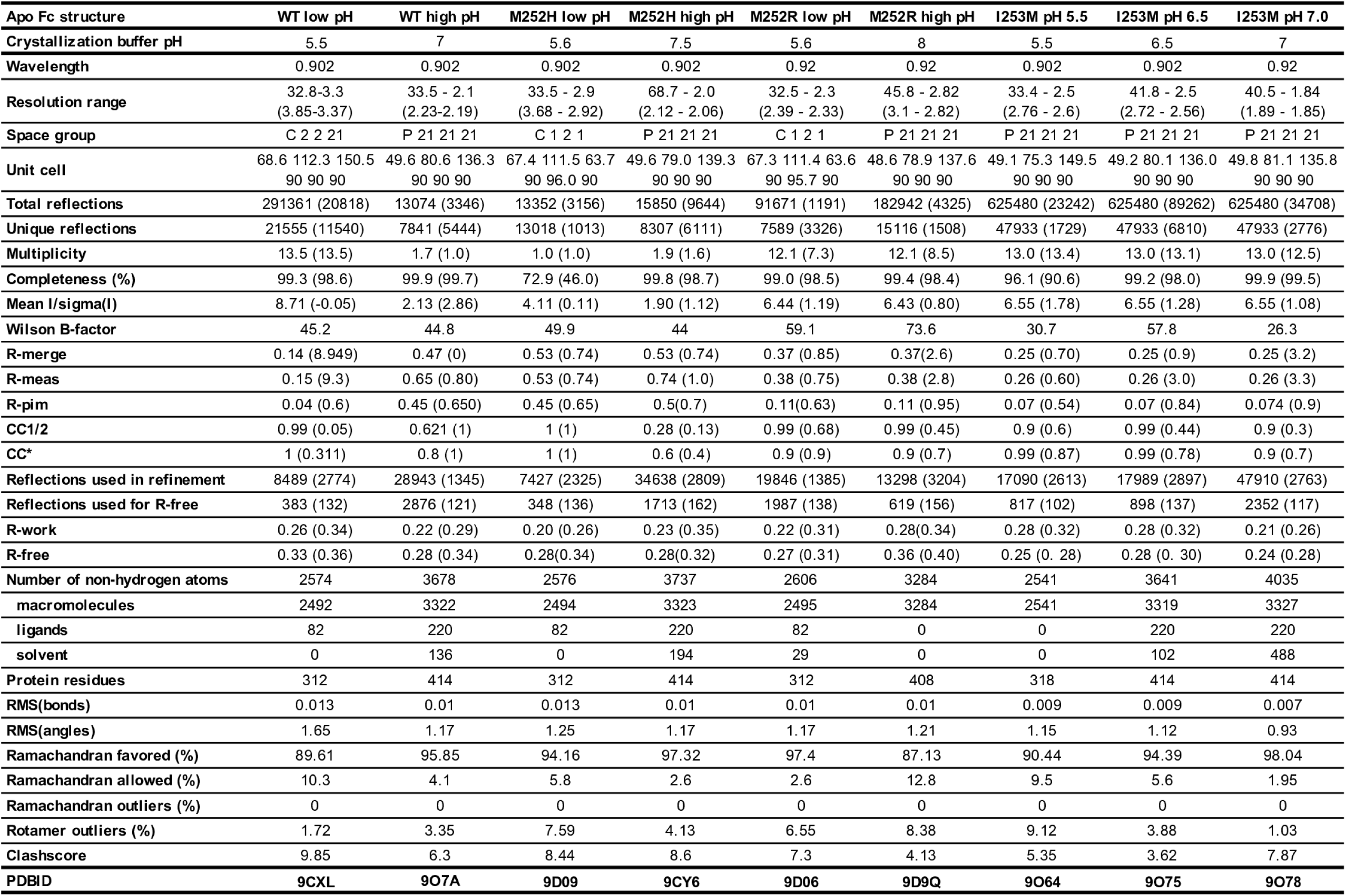
Crystallographic data collection and refinement statistics for apo Fc structures. All structures were solved by molecular replacement. Values in parentheses are for the highest resolution shell. Crystallization buffer pH and PDB Accession numbers are also shown.

**Supplemental Table 4.** Comparison of crystallographically solved structures from this study with three reference *wt* conformations. Structural differences are assessed using four metrics: RMSD’, RMSD”, angle, and displacement (detailed below). The structures (rows) are superimposed on the CH2 domain of the reference structures (columns). RMSD’ represents the root-mean-square deviation (RMSD) between CH2 domains, reflecting their structural similarity. From this aligned position, CH3 domain reorientation is quantified using displacement (translation in Å) and angle (rotation in °). RMSD” measures the RMSD between CH3 domains, capturing conformational divergence in this region. The reorientation of the CH3 domain was quantified using the https://pymolwiki.org/index.php/Angle_between_domains script in PyMOL.

| Reference structure | WT low pH (apo) | WT high pH (apo) | WT (holo) |
| --- | --- | --- | --- |
| WT low pH (apo) | - | (0.6, 0.5, 17.7, 4.6) | (0.6, 0.7, 14.8, 4.4) |
| M252H low pH (apo) | (0.4, 0.4, 2.3, 0.6) | (0.4, 0.4, 20.4, 5.6) | (0.4, 0.7, 16.8, 4.8) |
| M252R low pH (apo) | (0.4, 0.4, 2.4, 0.6) | (0.4, 0.3, 20.7, 6.0) | (0.4, 0.7, 16.8, 4.8) |
| WT high pH (apo) | (0.6, 0.5, 17.7, 4.6) | - | (0.4, 0.6, 10.1, 2.8) |
| M252H high pH (apo) | (0.6, 0.4, 17.6, 4.5) | (0.1, 0.2, 0.9, 0.4) | (0.4, 0.7, 9.0, 2.4) |
| M252R high pH (apo) | (0.6, 0.5, 12.8, 4.2) | (0.7, 0.3, 10.9, 4.2) | (0.5, 0.7, 2.1, 1.1) |
| WT holo (apo) | (0.6, 0.7, 14.8, 4.4) | (0.4, 0.6, 10.1, 2.8) | - |
| I253M pH=5.5 (apo) | (0.7, 0.6, 15.3, 3.7) | (0.4, 0.4, 4.7, 1.2) | (0.6, 0.7, 9.1, 2.1) |
| I253M pH=6.5 (apo) | (0.6, 0.5, 16.9, 4.2) | (0.2, 0.2, 0.9, 0.2) | (0.5, 0.7, 9.4, 2.8) |
| I253M pH=7.0 (apo) | (0.6, 0.4, 17.9, 4.6) | (0.1, 0.1, 0.5, 0.1) | (0.4, 0.6, 9.9, 2.8) |

**Supplemental Table 5.** Analysis of pairwise chemical interactions between the T307W residue and its neighboring residues in the solvent leg trajectories. To estimate the Δ*G* of the solvent leg, three independent 10 ns simulations were performed using the OPLS5 force field and different random seeds. The percent occurrence of each interaction was calculated using the analyze_trajectory_ppi.py script from the Schrödinger suite. Key interactions contributing to solvent leg energy differences (described in Supplemental Figure 5B, right panel) are highlighted in bold text.

| **ΔG holo solvent** | **E258-T307W**<br>**ss Hbond** | E258-T307W<br>water bridge | **K288-T307W**<br>**cation-pi** | K288-T307W<br>hydrophobic | **T256-T307W**<br>**hydrophobic** | **T256-T307W**<br>**ss Hbond** | V305-T307W<br>hydrophobic | N286-T307W<br>water bridge | N286-T307W<br>hydrophobic |
| --- | --- | --- | --- | --- | --- | --- | --- | --- | --- |
| repeat1 (-4.76) | 55.81% | 6.98% | 51.16% | 51.16% | 79.07% | 0.00% | 32.56% | 74.42% | 27.91% |
| repeat2 (-6.33) | **65.12%** | 4.65% | **97.67%** | 20.93% | **97.67%** | 0.00% | 0.00% | 88.37% | 0.00% |
| repeat3 (-5.65) | 51.16% | 13.95% | 81.40% | 27.91% | 88.37% | 0.00% | 9.30% | 69.77% | 4.65% |

| **ΔG apo solvent** | **E258-T307W**<br>**ss Hbond** | E258-T307W<br>water bridge | **K288-T307W**<br>**cation-pi** | K288-T307W<br>hydrophobic | **T256-T307W**<br>**hydrophobic** | **T256-T307W**<br>**ss Hbond** | V305-T307W<br>hydrophobic | N286-T307W<br>water bridge | N286-T307W<br>hydrophobic |
| --- | --- | --- | --- | --- | --- | --- | --- | --- | --- |
| repeat1 (-3.97) | 41.86% | 4.65% | 74.42% | 20.93% | 44.19% | 4.65% | 0.00% | 79.07% | 4.65% |
| repeat2 (-4.77) | 44.19% | 4.65% | 32.56% | 93.02% | 32.56% | 0.00% | 72.09% | 60.47% | 9.30% |
| repeat3 (-4.71) | 44.19% | 16.28% | 30.23% | 81.40% | 27.91% | 2.33% | 48.84% | 41.86% | 25.58% |

**Supplemental Table 6.** Dependence of solvent leg FEP energies on variations in the local environment near the mutation site in the input structures. Δ*G* values for the solvent leg were calculated from 10 ns simulations using the OPLS5 force field. Forward FEP simulations (Thr307 → Trp, Gln, neutral or charged Glu) were initiated from one of the three most common rotameric states of Thr in either the apo or holo conformation. Reverse simulations (Trp307 → Thr) used input geometries shown in Supplemental Figure 5C.

| **Forward**<br>**mutation** | apo<br>rotamer1 | apo<br>rotamer2 | apo<br>rotamer3 | holo<br>rotamer1 | holo<br>rotamer2 | holo<br>rotamer3 | Range |
| --- | --- | --- | --- | --- | --- | --- | --- |
| THR307TRP | -4.81 | -4.58 | -4.74 | -4.07 | **-5.25** | -4.07 | 1.18 |
| THR307GLN | -48.70 | -48.78 | -49.23 | -48.79 | -49.06 | -49.01 | 0.53 |
| THR307GLU | -156.02 | -155.81 | -155.63 | -156.19 | -155.81 | -156.07 | 0.56 |
| THR307GLH | -36.86 | -36.91 | -36.98 | -37.04 | -37.29 | -36.75 | 0.54 |

| **Reverse**<br>**mutation** | holo stab | holo<br>rotamer1 | holo<br>rotamer2 | holo<br>rotamer3 |
| --- | --- | --- | --- | --- |
| TRP307THR | **5.38** | 5.07 | 4.78 | 4.81 |
|  | apo stab | apo<br>rotamer1 | apo<br>rotamer2 | apo<br>rotamer3 |
|  | **5.61** | 4.47 | 5.46 | 4.55 |

**Supplemental Table 7.**
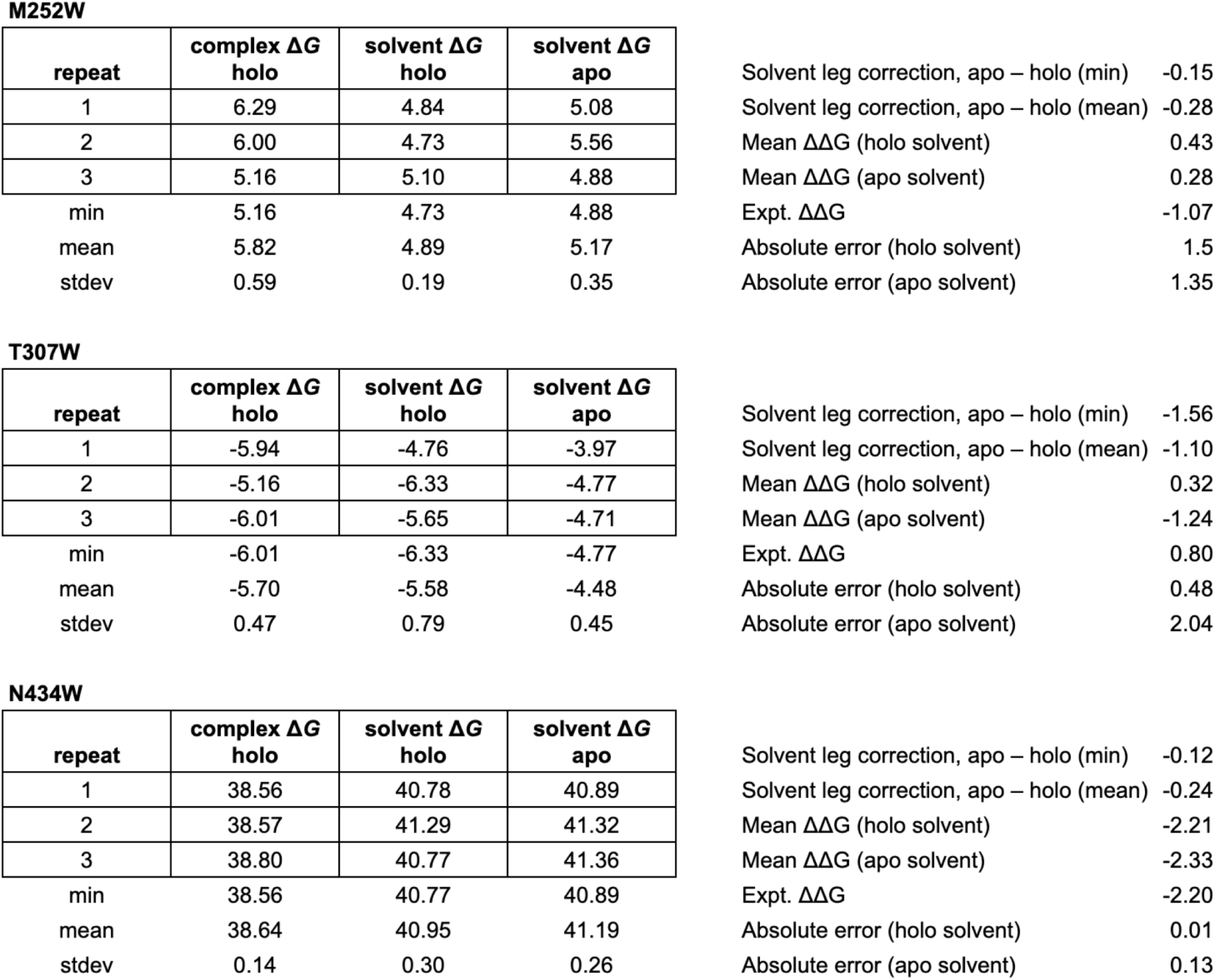
Comparison of solvent leg energies using holo and apo conformations, from 10 ns Protein FEP simulations using OPLS5 force field; all Δ*G*/ΔΔ*G* values in kcal/mol.

**M252W**
| repeat | complex $\Delta G$<br>holo | solvent $\Delta G$<br>holo | solvent $\Delta G$<br>apo |
| --- | --- | --- | --- |
| 1 | 6.29 | 4.84 | 5.08 |
| 2 | 6.00 | 4.73 | 5.56 |
| 3 | 5.16 | 5.10 | 4.88 |
| min | 5.16 | 4.73 | 4.88 |
| mean | 5.82 | 4.89 | 5.17 |
| stdev | 0.59 | 0.19 | 0.35 |

|  |  |
| --- | --- |
| Solvent leg correction, apo – holo (min) | -0.15 |
| Solvent leg correction, apo – holo (mean) | -0.28 |
| Mean $\Delta\Delta G$ (holo solvent) | 0.43 |
| Mean $\Delta\Delta G$ (apo solvent) | 0.28 |
| Expt. $\Delta\Delta G$ | -1.07 |
| Absolute error (holo solvent) | 1.5 |
| Absolute error (apo solvent) | 1.35 |

**T307W**
| repeat | complex $\Delta G$<br>holo | solvent $\Delta G$<br>holo | solvent $\Delta G$<br>apo |
| --- | --- | --- | --- |
| 1 | -5.94 | -4.76 | -3.97 |
| 2 | -5.16 | -6.33 | -4.77 |
| 3 | -6.01 | -5.65 | -4.71 |
| min | -6.01 | -6.33 | -4.77 |
| mean | -5.70 | -5.58 | -4.48 |
| stdev | 0.47 | 0.79 | 0.45 |

|  |  |
| --- | --- |
| Solvent leg correction, apo – holo (min) | -1.56 |
| Solvent leg correction, apo – holo (mean) | -1.10 |
| Mean $\Delta\Delta G$ (holo solvent) | 0.32 |
| Mean $\Delta\Delta G$ (apo solvent) | -1.24 |
| Expt. $\Delta\Delta G$ | 0.80 |
| Absolute error (holo solvent) | 0.48 |
| Absolute error (apo solvent) | 2.04 |

**N434W**
| repeat | complex $\Delta G$<br>holo | solvent $\Delta G$<br>holo | solvent $\Delta G$<br>apo |
| --- | --- | --- | --- |
| 1 | 38.56 | 40.78 | 40.89 |
| 2 | 38.57 | 41.29 | 41.32 |
| 3 | 38.80 | 40.77 | 41.36 |
| min | 38.56 | 40.77 | 40.89 |
| mean | 38.64 | 40.95 | 41.19 |
| stdev | 0.14 | 0.30 | 0.26 |

|  |  |
| --- | --- |
| Solvent leg correction, apo – holo (min) | -0.12 |
| Solvent leg correction, apo – holo (mean) | -0.24 |
| Mean $\Delta\Delta G$ (holo solvent) | -2.21 |
| Mean $\Delta\Delta G$ (apo solvent) | -2.33 |
| Expt. $\Delta\Delta G$ | -2.20 |
| Absolute error (holo solvent) | 0.01 |
| Absolute error (apo solvent) | 0.13 |

### Supplementary Figures

**Supplemental Figure 1.**
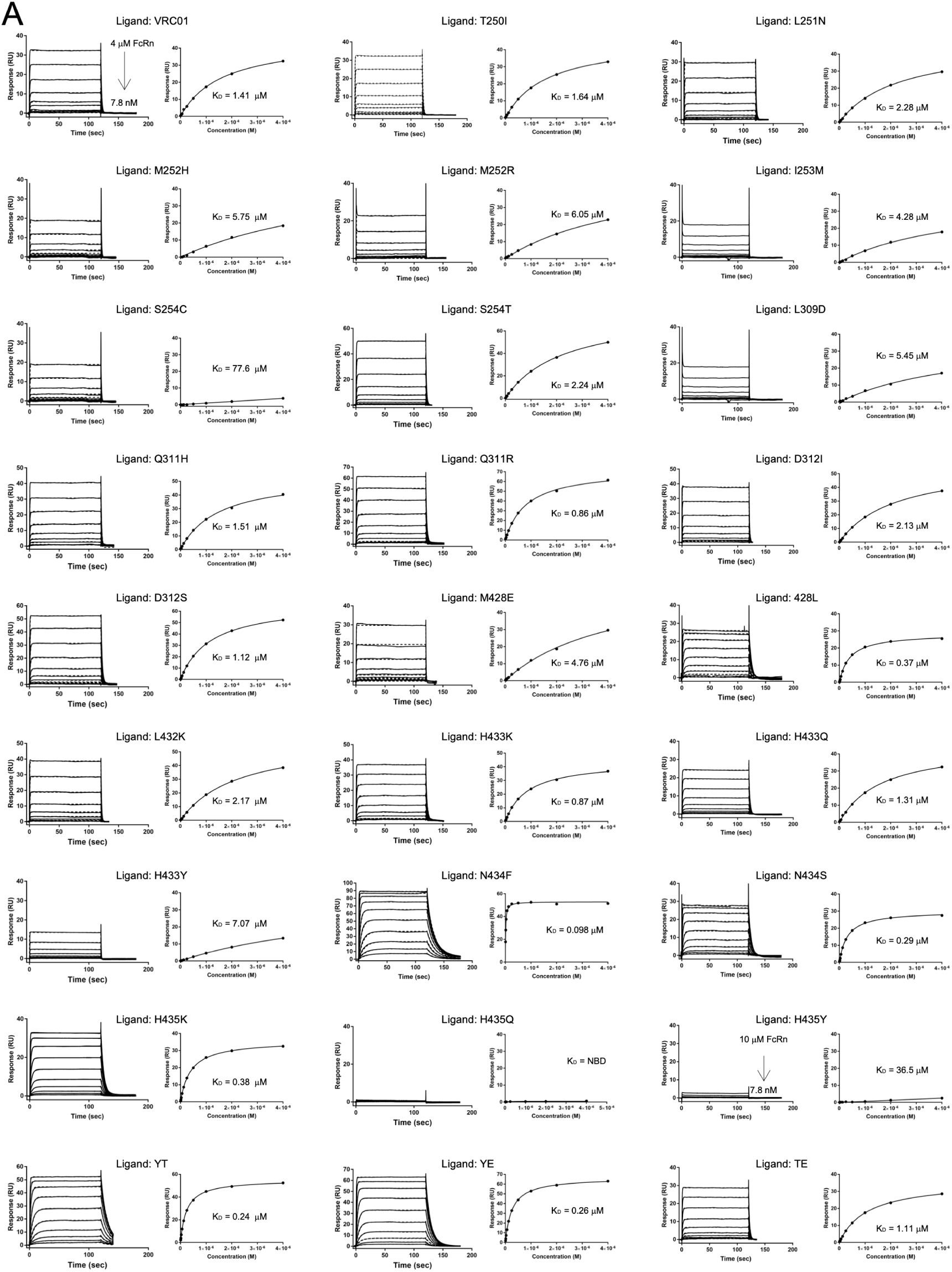
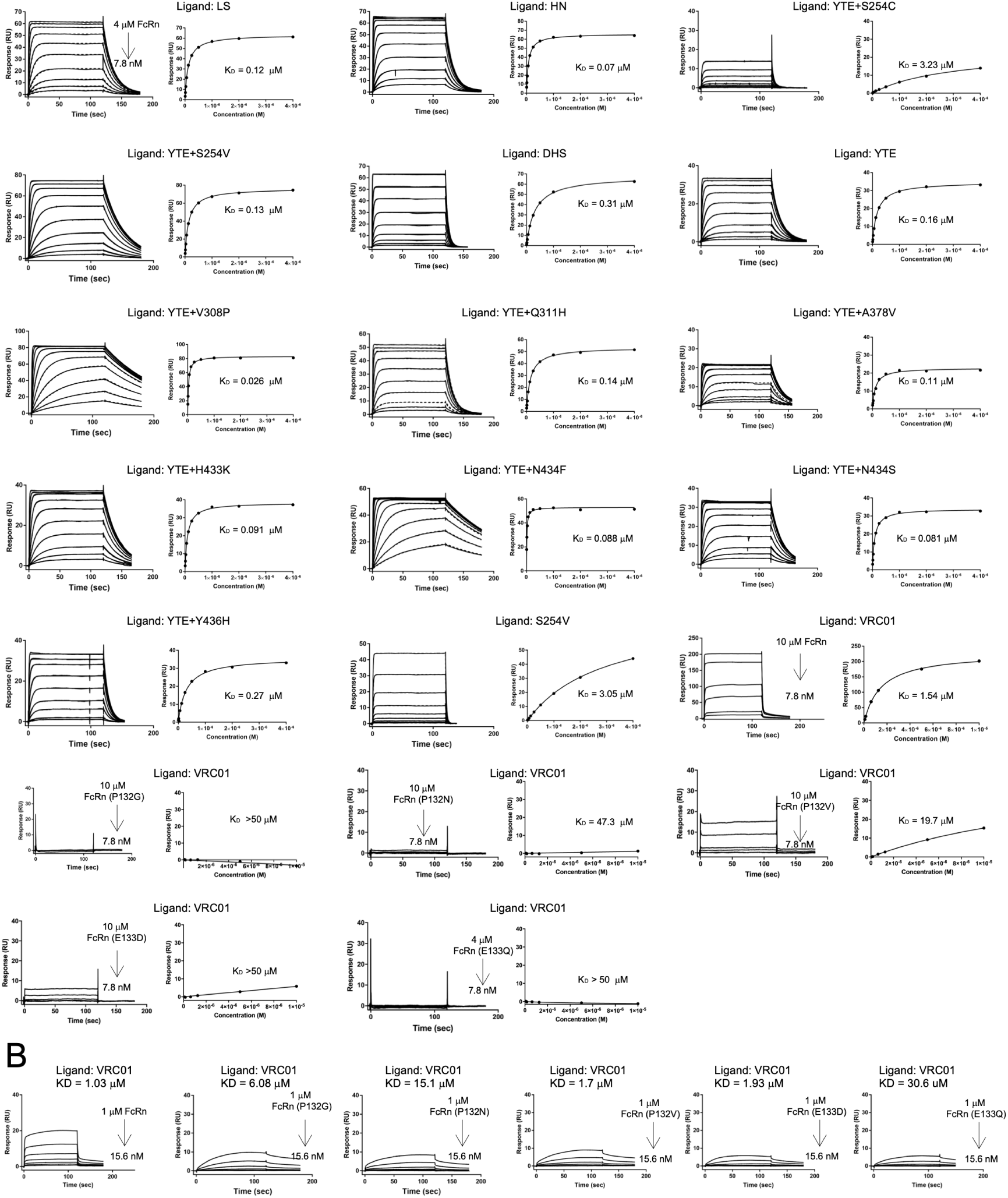
SPR binding measurements for Fc-FcRn variants at pH 6.0. (A) *Left panels:* Binding sensorgrams for concentration series of *wt* FcRn analyte injected over immobilized Fc-variant VRC01 ligands. *Right panels:* Equilibrium binding responses plotted on a linear scale against injected FcRn variant concentrations and fit to a simple binding isotherm. Inset *K*_D_ values correspond to the FcRn concentration that generated 0.5 R_max_. (B) Binding sensorgrams for variant FcRn analytes injected over immobilized *wt* VRC01 ligand, and fit to a 1:1 Langmuir binding model. Inset *K*_D_ values correspond to the ratio between dissociation and association rate constants, 𝑘_off_ /𝑘_on_.

**Supplemental Figure 2.**
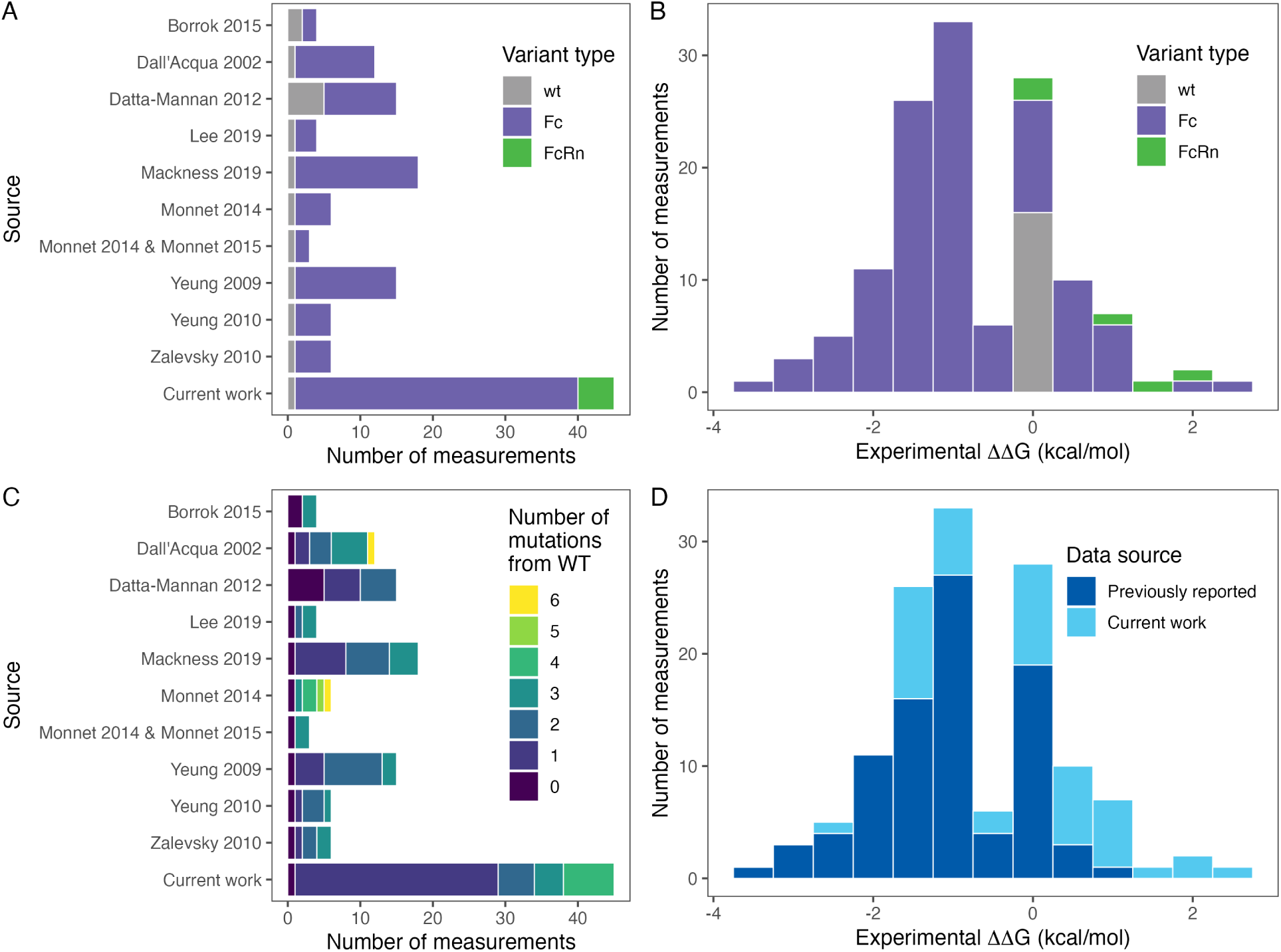
Histograms providing additional information about the experimental dataset. (A) Number of experimental Fc-FcRn variant binding affinity measurements obtained from each published source and the current work, colored by mutated protein. (B) Distribution of experimental binding free energies (ΔΔ*G*) of the full dataset relative to the *wt* Fc-FcRn complex, colored as in (A). (C) Breakdown of the number of mutations from *wt* Fc-FcRn present in the variants from each source. (D) Distribution of experimental ΔΔ*G*s, showing increased dynamic range and improved coverage of postive ΔΔ*G* values after including the measurements from the current work.

**Supplemental Figure 3.**
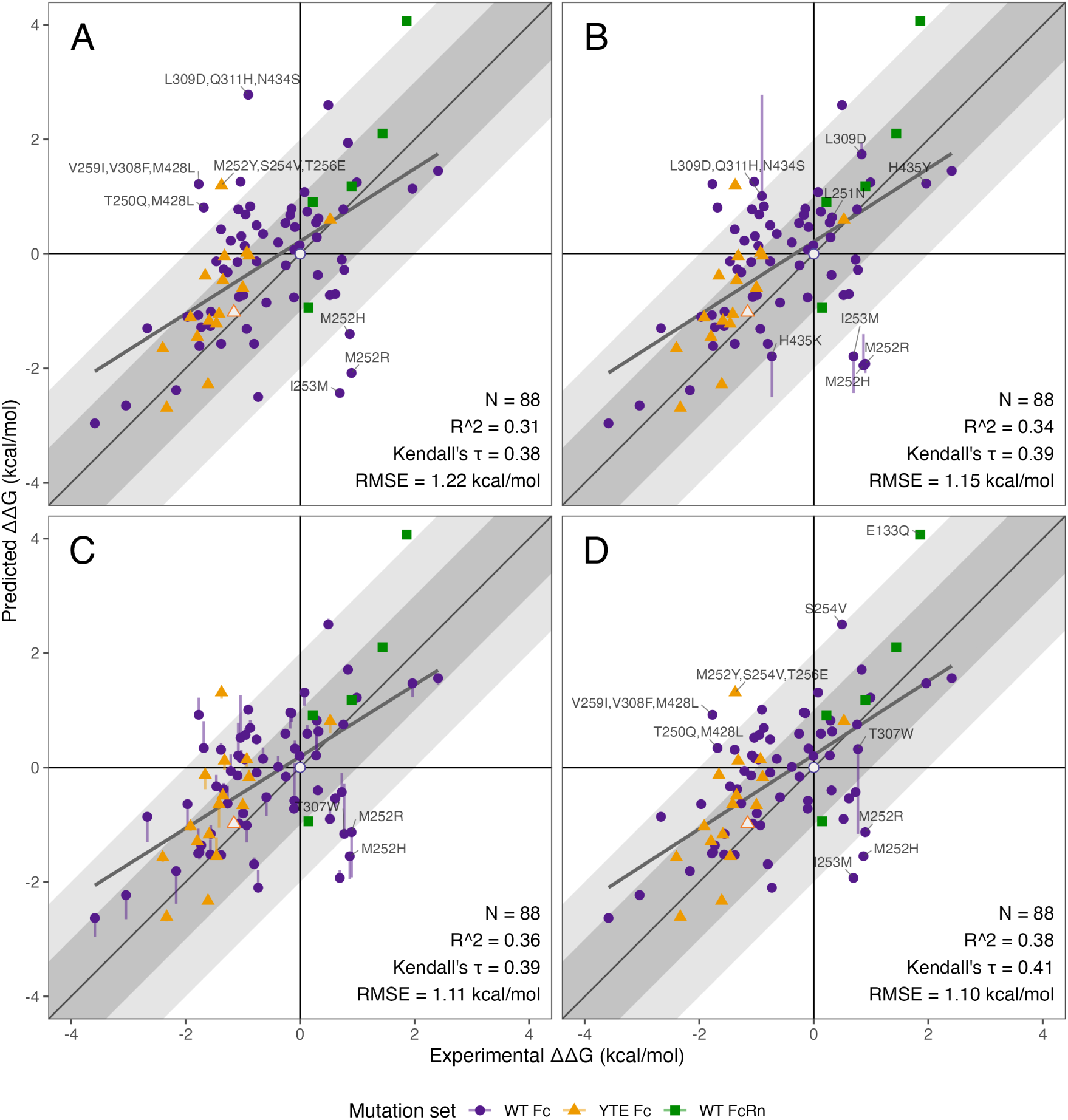
Intermediate retrospective FEP results. Results are shown from 10 ns FEP simulations at four stages: (A) original results using OPLS5 force field and default protocols; (B) accounting for coupling between mutation sites and key titratable Fc His residues H310 and H435; (C) incorporating apo Fc solvent leg Δ*G* values; and (D) addressing a local conformational difference in the apo model for the Fc T307W variant. Vertical tails indicate change in predicted ΔΔ*G* from the previous stage.

**Supplemental Figure 4.**
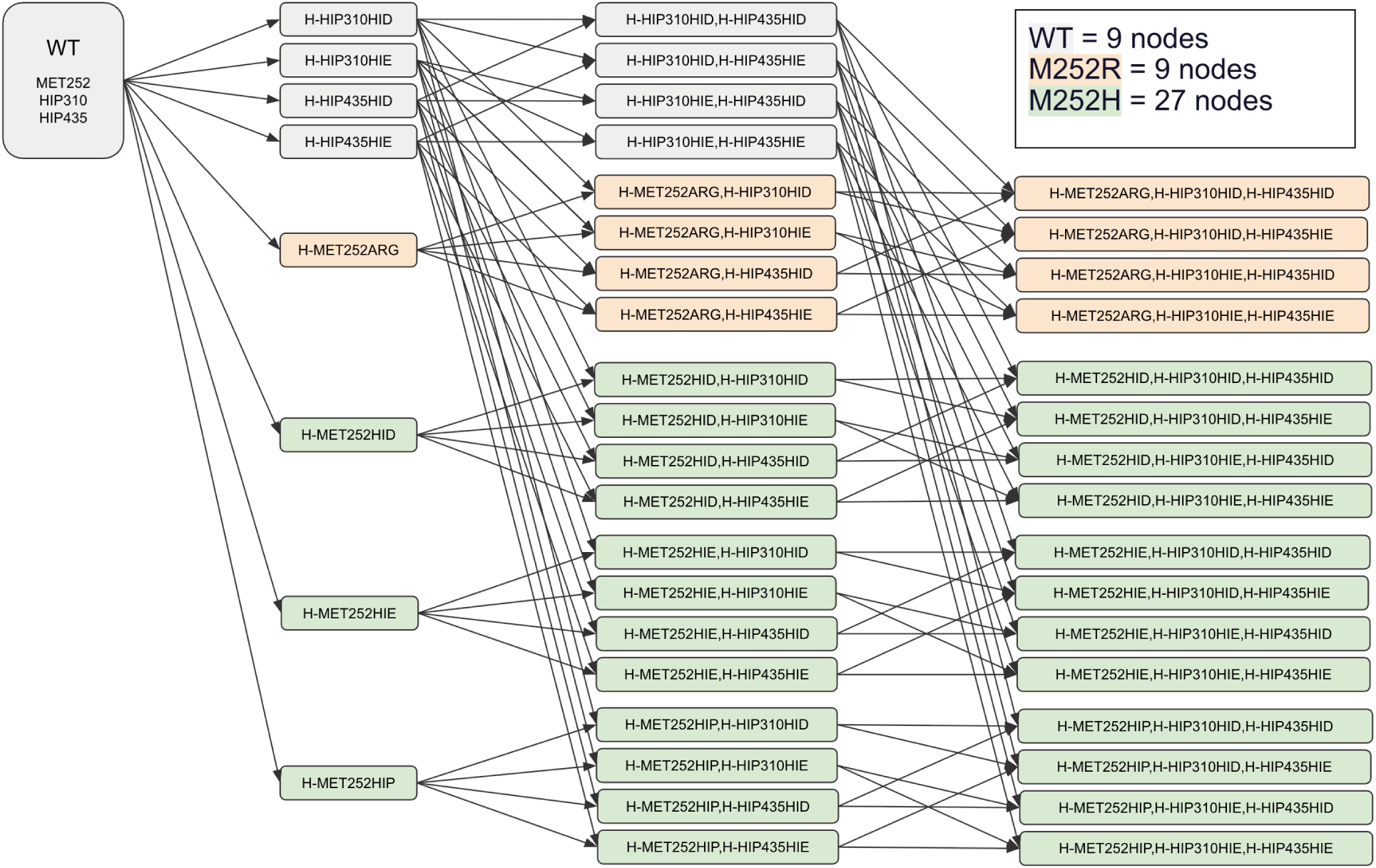
Example illustration of Protein FEP graph topology for Fc M252R and M252H variants with explicit coupled titration of Fc H310 and H435.

**Supplemental Figure 5.**
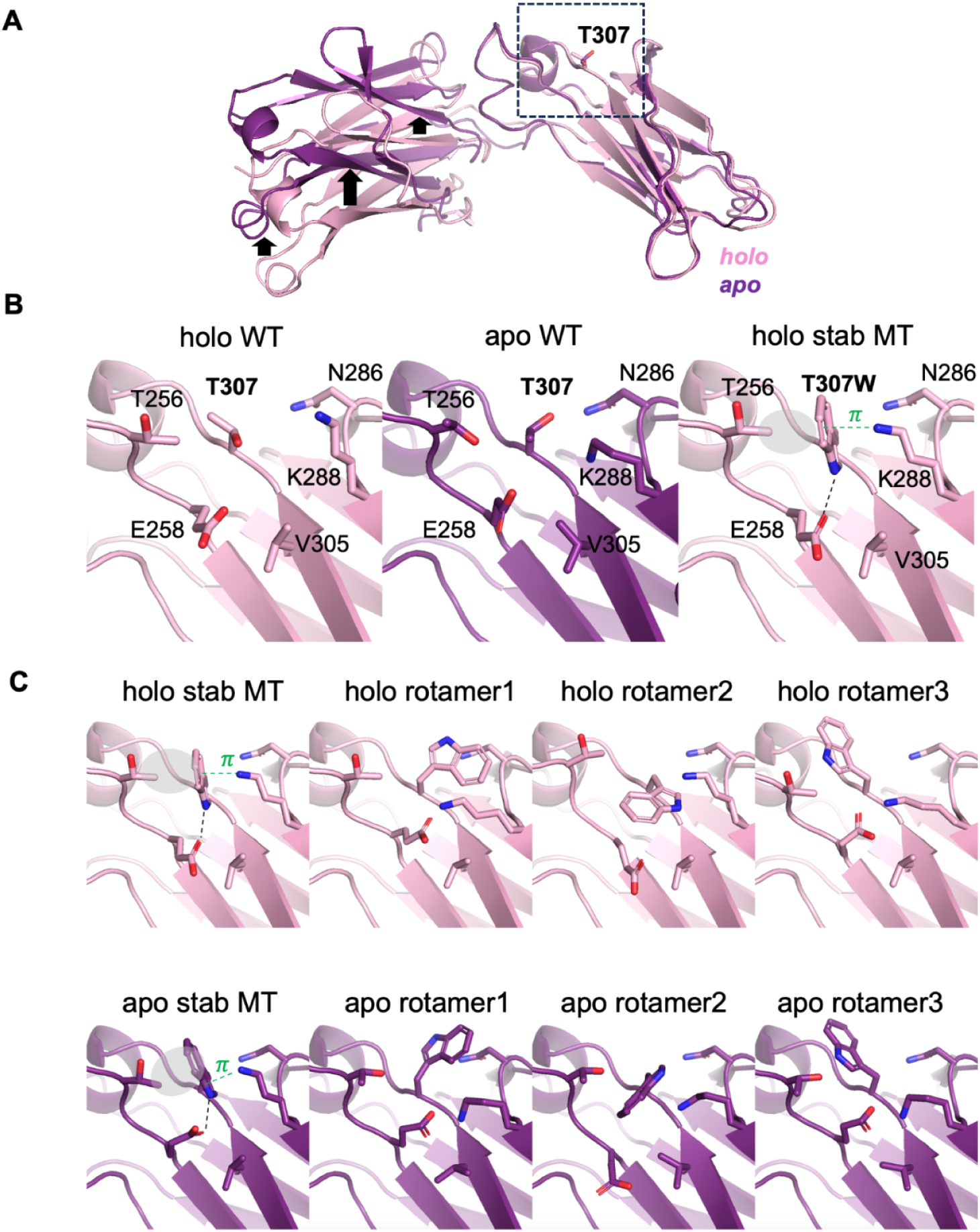
Structural heterogeneity contributing to convergence issues in the T307W outlier. (A) Ribbon representation of holo (pink) and apo (purple) Fc conformations aligned on the CH2 domain, with the mutation site indicated by a dashed box, far from the region undergoing conformational change. (B) Close-up view of the boxed region in (A), showing the side chain of residue 307 and surrounding amino acids in stick representation. The stable interaction network observed in simulations with low energy of the mutant state (right panel) includes a hydrophobic contact between T256 and T307W (gray oval), a cation-π interaction between K288 and T307W (green), and a hydrogen bond between E258 and T307W. (C) Input structures used for reverse FEP simulations (Trp307→Thr).

**Supplemental Figure 6.**
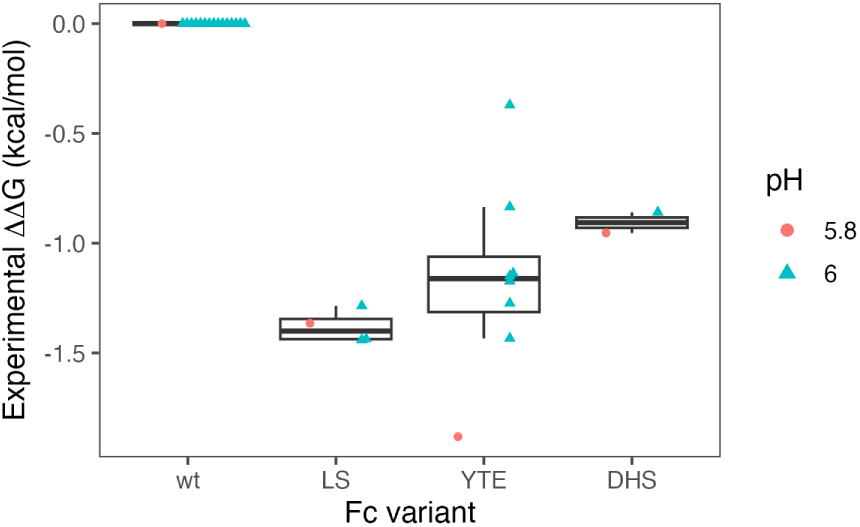
Experimental ΔΔ*G* values relative to *wt* for pH 5.8 Fc variant binding affinity measurements from Lee, et al. [14] compared to pH 6.0 ΔΔ*G* values for the same variants from other sources.

